# Backward masking reveals coarse-to-fine dynamics in human V1

**DOI:** 10.1101/2023.02.08.525486

**Authors:** Jolien P. Schuurmans, Matthew A. Bennett, Kirsten Petras, Valérie Goffaux

## Abstract

Natural images exhibit luminance variations aligned across a broad spectrum of spatial frequencies (SFs). It has been proposed that, at early stages of processing, the coarse signals carried by the low SF (LSF) of the visual input are sent rapidly from primary visual cortex (V1) to ventral, dorsal and frontal regions to form a coarse representation of the input, which is later sent back to V1 to guide the processing of fine-grained high SFs (i.e., HSF). We used functional resonance imaging (fMRI) to investigate the role of human V1 in the coarse-to-fine integration of visual input. We disrupted the processing of the coarse and fine content of full-spectrum human face stimuli via backward masking of selective SF ranges (LSFs: <1.75cpd and HSFs: >1.75cpd) at specific times (50, 83, 100 or 150ms). In line with coarse-to-fine proposals, we found that (1) the selective masking of stimulus LSF disrupted V1 activity in the earliest time window, and progressively decreased in influence, while (2) an opposite trend was observed for the masking of a stimulus’ HSF. This pattern of activity was found in V1, as well as in ventral (i.e. the Fusiform Face area, FFA), dorsal and orbitofrontal regions. We additionally presented participants with contrast negated stimuli. While contrast negation significantly reduced response amplitudes in the FFA, as well as coupling between FFA and V1, coarse-to-fine dynamics were not affected by this manipulation. The fact that V1 response dynamics to strictly identical stimulus sets differed depending on the masked scale adds to growing evidence that V1 role goes beyond the early and quasi-passive transmission of visual information to the rest of the brain. It instead indicates that V1 may yield a ‘spatially registered common forum’ or ‘blackboard’ that integrates top-down inferences with incoming visual signals through its recurrent interaction with high-level regions located in the inferotemporal, dorsal and frontal regions.

## Introduction

Visual perception results from the recurrent (i.e., iterative) communication within and across different levels of the cortical hierarchy via horizontal, feedforward, and feedback connections (e.g., Gilbert & Li, 2013; Hupe et al., 1998; Kravitz et al., 2013; Kreiman & Serre, 2020; Lamme et al., 1998; Petro et al., 2014). How recurrent processing converts the incoming visual input into a stable and rich perceptual experience, is still not completely understood (Olshausen & Field, 2006; Roland, 2010). One line of inquiry stems from the notion that visual function, therefore recurrent processing, has adapted to regularities in the spatial structure of the visual environment (over onto- and phylogeny; Barlow, 1975). One fundamental regularity is that natural images contain luminance variations over a wide spectrum of spatial scales, or frequencies (SFs) and that, across SFs, such luminance variations align in phase. Several authors proposed that recurrent processing exploits SF alignment to support the efficient integration of visual information (Bar, 2004; Bullier, 2001; Hegde, 2008; Watt, 1987; Kreiman & Serre, 2020). In this framework, visual processing starts with the fast feedforward transmission of the coarse structure of the input conveyed by low SFs (LSF), presumably via the rapid and transient magnocellular pathway. Within the first tens of milliseconds of processing, based on this initial sweep a coarse representation of the visual input is generated in high-level visual regions that retain prior knowledge about the natural statistics of the category best matching input (e.g., Bar, 2004; Lee, 2015; Mumford, 1992). These high-level coarse representations in dorsal/parietal, frontal and ventral cortex are then fed back to V1 (and possibly V2) and used to guide the processing of the fine-grained cues that are contained in high SF (HSF) input relayed via the slower and more sustained parvocellular pathway (Baseler & Sutter, 1997; Nowak et al., 1995; Schmolesky et al., 1998). V1 (and V2) is proposed to operate as the ‘active blackboard’ of the coarse-to-fine integration of visual input. There are several hints that V1 yields such a ‘spatially registered common forum’ or ‘blackboard’ (Bullier, 2001; Deco & Lee, 2004; Roland, 2010) able to receive and integrate higher-order inferences with incoming visual signals via recurrent interactions from the rest of the brain. First, V1 encodes visual input with the greatest spatial specificity; it is thus necessarily involved for the sharp representation of the visual input (Bullier, 2001; Deco & Lee, 2004; Lee et al., 1998). Moreover, V1 neurons attune to progressively higher SFs over the course of processing in line with a core tenet of coarse-to-fine proposals, namely that LSFs are encoded before HSFs (Allen & Freeman, 2006; Bredfeldt & Ringach, 2002; Einevoll et al., 2011; Frazor et al., 2004; Malone et al., 2007; Mazer et al., 2002; Nirody, 2014; Parker & Salzen, 1977; Tanaka & Sawada, 2022). Lastly, the fact that V1 is highly interconnected with the rest of the brain, with thalamic afferences thought to account for a very limited fraction of V1 activity (Budd, 1998; Douglas & Martin, 2007; Kravitz et al., 2013; Markov et al., 2014; Muckli & Petro, 2013), indicates that responses in this region are largely modulated by non-retinal signals.

Despite the increasing influence of coarse-to-fine theories in vision and computer sciences (for a review see e.g. Kreiman & Serre, 2020), and the fact that V1 is the best documented region in the primate brain, evidence for its contribution to human vision as an ‘active blackboard’ or ‘high-resolution buffer’ for coarse-to-fine integration is surprisingly scarce. Past research supports the notion that coarse input scales are processed before fine details (for simple stimuli such as lines, dots and gratings see e.g. Breitmeyer (1975); Hughes et al. (1996); Jones and Keck (1978); Mihaylova et al. (1999); Musselwhite and Jeffreys (1985); Parker (1980); Parker and Dutch (1987); Watt (1987); and for complex stimuli such as faces and natural scenes see e.g. Goffaux et al. (2011); Halit et al. (2006); Kauffmann et al. (2015b); Kauffmann et al. (2015d); Lu et al. (2018); Musel et al. (2014); Parker et al. (1992, 1997); Peyrin et al. (2006); Schyns and Oliva (1994); Vlamings et al. (2009)). However, because this research investigates LSF and HSF processing separately, via the use of SF-filtered stimuli, it cannot address a fundamental prediction of recurrent processing, i.e. that coarse-to-fine integration helps the visual processing of natural, full-spectrum, input. We recently addressed this question and showed that the availability of meaningful LSF information, as found in broadband phase-aligned images, indeed reduces the resources allocated to HSF processing in the (extra)striate cortex, compared to phase-scrambled images (Petras et al., 2021; Petras et al., 2019). This suggests that, in natural (full-spectrum) visual input where SFs align in phase and thus contain partially redundant information, the rapidly processed LSF information guides the processing of the rich and complex HSF, therefore reducing computational costs. Yet, the spatial resolution afforded by MEG (Petras et al., 2021; or EEG as in Petras et al., 2019) was insufficient to discern whether V1 itself is involved in the coarse-to-fine integration.

Here we systematically addressed the contribution of V1 to the coarse-to-fine integration of visual information. We reasoned that if coarse-to-fine integration is implemented in V1 for the sake of efficient processing, its disruption should induce a measurable cost in V1 processing (i.e., less sharp/sparse representations), which, in this region, typically results in an enhanced level of Blood-Oxygen level-dependent, i.e., BOLD, response (e.g., de-Wit et al., 2012; Freud et al., 2017; Goffaux et al., 2016; Kok & de Lange, 2014; Markov et al., 2014; Olshausen & Field, 1996; but see Rieger et al., 2013). We disrupted the coarse-to-fine recurrent processing of human face stimuli via backward masking (Bacon-Mace et al., 2005; Breitmeyer, 1984; Fahrenfort et al., 2007; Grill-Spector et al., 2000; Kovacs et al., 1995; Lamme et al., 2002; Rolls & Tovee, 1994; Rolls et al., 1999). We showed full-spectrum face images for either 50, 83, 100 or 150ms; these stimulus onset asynchronies (SOAs) were chosen to track recurrent processing from its early stage on (see Ahmed et al., 2008; Eriksson & Roland, 2006; Harvey et al., 2009; Lamme, 1995; Roland, 2010; Roland et al., 2006; Scholte et al., 2008; Takagaki et al., 2008; Wibral et al., 2009). The mask contained either LSF or HSF noise, to selectively disrupt the coarse or fine processing of the face stimulus (Bacon-Mace et al., 2005; Gao & Bentin, 2011; Stromeyer & Julesz, 1972). This approach allowed us to disentangle the respective SF contribution to neural processing over time while accommodating a full range of aligned SFs in the stimuli (a critical aspect of naturalistic vision for the coarse-to-fine framework). We compared the backward masked responses between phase-aligned and phase-scrambled face stimuli in order to extract the visual responses specific to the processing of face shapes. In unconstrained viewing conditions, V1 activity increases in response to stimulus scrambling (Freud et al., 2017; Goffaux et al., 2016; Lerner et al., 2001), resulting in an overall negative intact-scrambled difference presumably due to an increase in the resources allocated to HSF processing (see Petras et al., 2021; Petras et al., 2019). Following coarse-to-fine theoretical proposals, interrupting LSF processing early prevents top-down LSF-driven signals from guiding the formation of a sparse representation, therefore increasing V1 responses to the face stimulus and resulting in an almost null intact-scrambled difference. In line with the presumably decreasing contribution of coarse scales over time, LSF masking should cause progressively less disturbance over time. This is expected to result in a progressive decrease of the intact-scrambled differential response, until negative values are reached, indexing the successful formation of a face representation in V1. We anticipate the reverse tendency with the masking of HSF as the contribution of HSF processing to perception is hypothesized to increase over processing course. This cross-like pattern of SF masking effects is the expected ‘signature’ of coarse-to-fine processing in V1. The high-level fusiform, frontal and parietal regions presumed to interact iteratively with V1 during coarse-to-fine integration were also expected to show a cross-like pattern. Since these regions increase their response as a function of the input visibility (Kay & Yeatman, 2016; Yue et al., 2013), we expected an overall increase of their response as a function of SOA, irrespective of the masked SF. Still, we expected such increase to be steeper for HSF masking. The present design imposes a stringent test of the coarse-to-fine predictions as the stimuli are full-spectrum and identical across masking conditions so that any difference reflects the differential contribution of SF to vision over processing course. The coarse representation initially formed in high-level visual regions has been proposed to involve some prior knowledge about the natural statistics of the category best matching input (e.g., Bar, 2004; Lee, 2015; Mumford, 1992). We thus expected contrast negation to hamper coarse-to-fine processing in V1 as this simple and reversible image manipulation disrupts the natural statistics of the human face stimulus (George et al., 1999; Liu-Shuang et al., 2022; Yue et al., 2013).

## Methods

### Subjects

Eighteen healthy subjects (age 25.1 ± 3.28, 5 males) - with no history of neurological disease - completed three scanning sessions each. Two subjects were excluded from data analysis due to inattentiveness and motion in the scanner. Before scanning, we tested the visual acuity of the subjects with the Freiburg Visual Acuity and Contrast Test (FrACT), which revealed all subjects had normal or corrected-to-normal vision (LogMar mean = -0.01 ± 0.11, decVA = 1.08 ± 0.39). All subjects reported being right-handed as measured with the Edinburgh handedness Questionnaire (82.9 ± 17.3; Oldfield, 1971). Subjects were compensated financially, and a signed informed consent was obtained. The study was approved by the Ethical Committee of the Université Catholique de Louvain (CEHF 2016/13SEP/393).

### Stimuli

Each subject performed the main experiment as well as a functional localiser to delineate the face-specialised voxels in the ventral pathway at the individual level. For both the funcitonal localiser and main experiment, the stimuli were presented with PsychoPy v3.2.4 on a NordicNeuroLab LCD Monitor (32-inch, 1920 × 1080 pixels, 698.40 × 392.85 mm, frame rate = 60 Hz) situated at the end of the scanner bore. Subjects viewed the stimuli via a mirror attached to the head-coil, at a viewing distance of 175cm. Across experiments, all images were attributed a mean (i.e., the global luminance) and root mean square (RMS) contrast of 0.45 ± 0.1.

#### Main experiment

We generated the main experimental stimuli based on 20 full-front greyscale photographs of unfamiliar human white faces (half male) with a neutral expression. Models were young adult alumni students (aged 18–25 years) of the Université Catholique de Louvain (Belgium), who gave written consent for the usage of their image (Laguesse et al., 2012). All faces were 367 pixels in height and 267 pixels in width and subtended a visual angle of 4.37° x 3.18°. To avoid the potential clipping of extreme intensity values in the image, we rescaled its luminance so that pixel values were between 0.1 and 0.9. Image luminance was then normalised (mean of zero and RMS contrast of 1). Next, faces were centred on a uniform grey background of 550 × 550 pixels (6.54° x 6.54° of visual angle).

Each face appeared in in three variations: natural, contrast negated, and phase-scrambled. Negated faces were created by subtracting the face pixel values from 1. We displayed the faces in a phase-scrambled version as a control condition. Phase scrambling preserves the amplitude spectrum of the original image but disrupts shape and edge information and hence image recognizability. We scrambled images by randomising their phase structure in the Fourier domain using a custom MATLAB function. Since manipulations in the Fourier domain operate on the whole image (face and background), the presence of a uniform background artificially increases the LSF energy of scrambled images. In order to minimise contamination of the phase-scrambled image by its originally uniform background, we employed an iterative phase scrambling procedure (as in Jacobs et al., 2020; Petras et al., 2021; Petras et al., 2019; see Figure 1a). Iterative scrambling involves first generating a phase scrambled version of the face image, then superimposing the original face back onto the scrambled image, scrambling this new image again, and repeating the procedure several times (for all our scrambled stimuli, 500 iterations were used). The iterative scrambling procedure results in an image in which the background and face share similar spectral properties. We employed iterative phase scrambling to create scrambled copies of the face stimuli (n=20), masks (n=40), and stimulus backgrounds (n=10).

**Figure 1.**
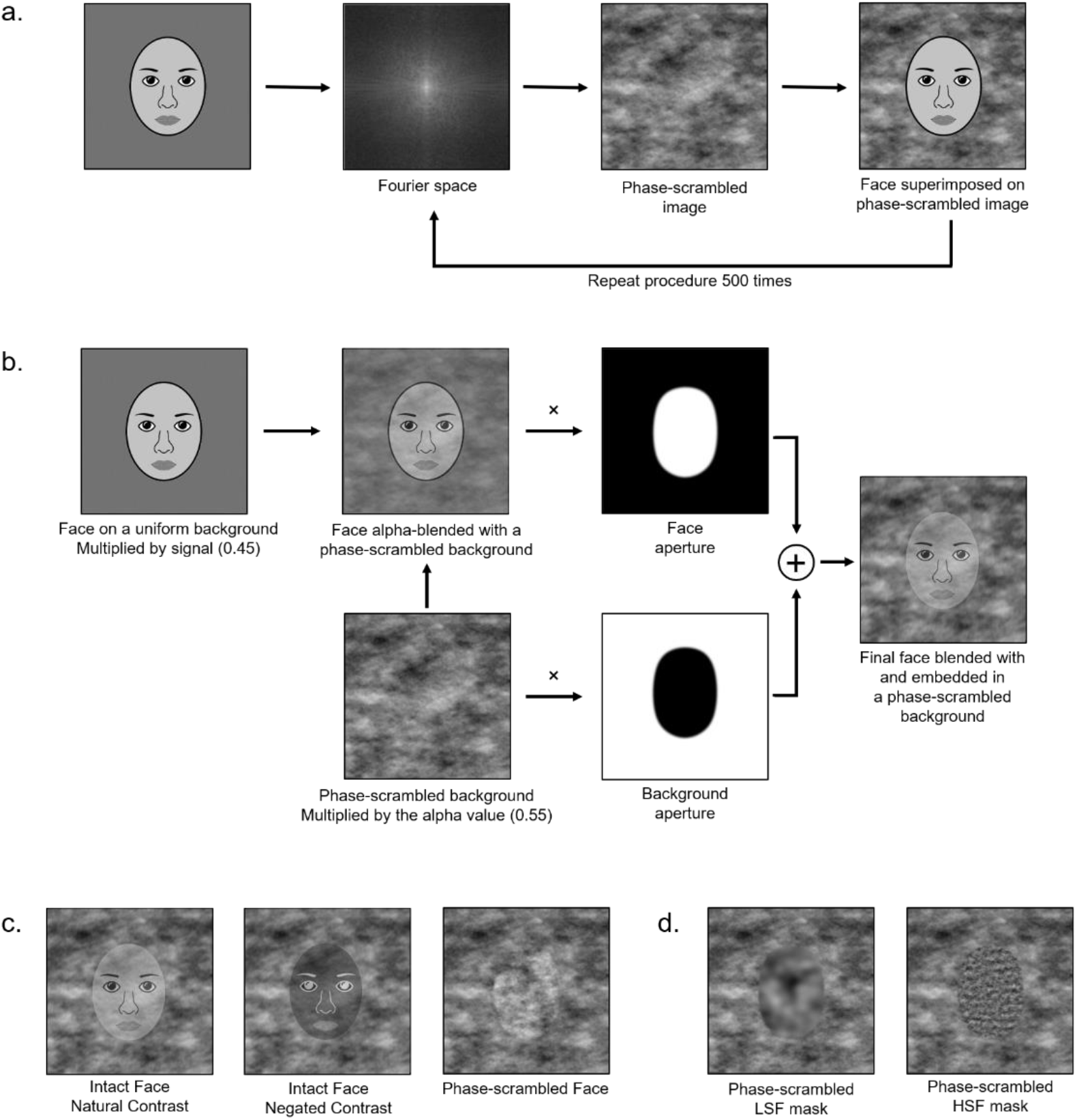
Stimulus generation. a) Iterative scrambling procedure. The image was first phase-scrambled in the Fourier domain, where after it was pasted back on the phase-scrambled image. This procedure was repeated 500 times until the final face-scrambled image had similar spectral properties as the face. b) To blend the face with the phase-scrambled background, we used alpha blending – an image combining method that relies on pixel transparency. Both the face image and background were multiplied with a face shaped aperture with a gradually transitioning border. The final face was embedded in and blended with the face scrambled background. c) All image types of an example face: intact natural and contrast negated stimuli, a phase-scrambled full-spectrum stimulus. The alpha blending was adapted for illustration purposes; experiment stimuli were less visible. d) Phase-scrambled masks containing either LSFs or HSFs. The masks were not alpha blended. Cartoon faces are used in replacement of photographs in the figure as journal regulations prohibit the use of identifiable faces.

We multiplied all stimuli and masks with a face-shaped aperture. Borders of this aperture gradually transitioned from 1 to 0 to avoid edge artefacts. A complementary aperture was multiplied with the scrambled background (see below) to avoid an increase of contrast in the transition region. To avoid abrupt stimulus on- and offset and instead provide a smooth transition between trial events within a block, all stimuli in a block were alpha-blended with a stable phase-scrambled background (e.g. see Ales et al., 2012; Figure 1b). Alpha blending creates a weighted sum of the stimulus and scrambled background contrast (a weight of 0.45 for stimulus and 0.55 for background) while maintaining mean luminance. The moderate RMS contrast of the stimuli and their reduced weight in the final alpha-blended stimulus were chosen to create challenging viewing conditions, and thus stimulate recurrent processing (Hupe et al., 1998; Kar & DiCarlo, 2021; Kar et al., 2019; Mohsenzadeh et al., 2018; Wyatte et al., 2014). As a result, the generated stimuli consisted of a face embedded in the phase-scrambled background (see Figure 1c). There were 20 blocks per condition, with the 10 phase-scrambled backgrounds repeated twice.

In contrast to the stimuli, the masks were superimposed on the background without alpha blending to ensure strong masking of the SF range it contains. Each stimulus (natural, negated, or scrambled face), and was then masked with a phase scrambled mask (Figure 1d). The scrambled masks were filtered to selectively contain either LSF or HSF, in order to selectively interfere with the processing of the LSF or HSF content of the full-spectrum face, respectively. In order to restrict mask image content to either LSFs or HSFs, we used 2nd order Butterworth filters with cut-offs such that four octaves were preserved for both SFs, without any gap between the two SF cut-offs (LSF: 0.7168 - 11.47cpi, HSF: 11.47 - 183.5cpi).

#### Functional localiser

The stimuli used for the localiser experiment were adapted from prior studies, available through the *fLoc functional localizer package* (Stigliani et al., 2015). Each of the three image categories - faces, hands, and instruments - contained 20 greyscale images (Supplementary material 1) as well as their phase-scrambled counterparts. The stimuli consisted of unfamiliar faces (half male) of various viewpoints (including hair), isolated hands in various poses, and stringed instruments (e.g. guitar, cello, lute, etc) positioned in different orientations. They were superimposed onto Fourier phase-scrambled backgrounds using the iterative scrambling procedure described above.

### Procedure

All subjects participated in 20 experimental runs divided over three scanning sessions. The first session included a T1-weighted anatomical scan and the second session a functional face localising run. Experimental and localiser functional runs all had a block design, of 10-second blocks, alternated by 10 seconds of fixation with a 12-second fixation period at the beginning and end of the run. A fixation period consisted of a uniform grey screen. Throughout all runs, a fixation cross was visible, consisting of two thin black lines connecting opposite corners of the square stimuli. Subjects were instructed to fixate on its central intersection at all times.

To ensure subjects paid attention during functional runs, we instructed them to detect a rare and brief colour change of the stimulus by pressing a button with the right index finger. In each block, there were two targets (red for the main experimental runs: HSV profile = 0.8, 1.0, 1.0, blue for the functional localiser run: HSV profile 1.0, 1.0, 0.8). One colour change occurred per block half, but never during the first stimulus of a block.

#### Main experiment

In each 8.7-minute run (see Figure 2a), there were 24 experimental conditions and two retinotopic localiser conditions. The order of experimental blocks was randomised with the constraint that no condition followed any other particular condition more than three times during the course of the experiment. The retinotopic localiser conditions at the end of every run consisted of a single block of 4Hz contrast-reversing checkerboard stimuli: in one condition limited to the region where the face appeared in the experimental conditions and the other to the background. This was used to localise the region of the (extra)striate cortex responding to the stimulus region containing the face selectively.

**Figure 2.**
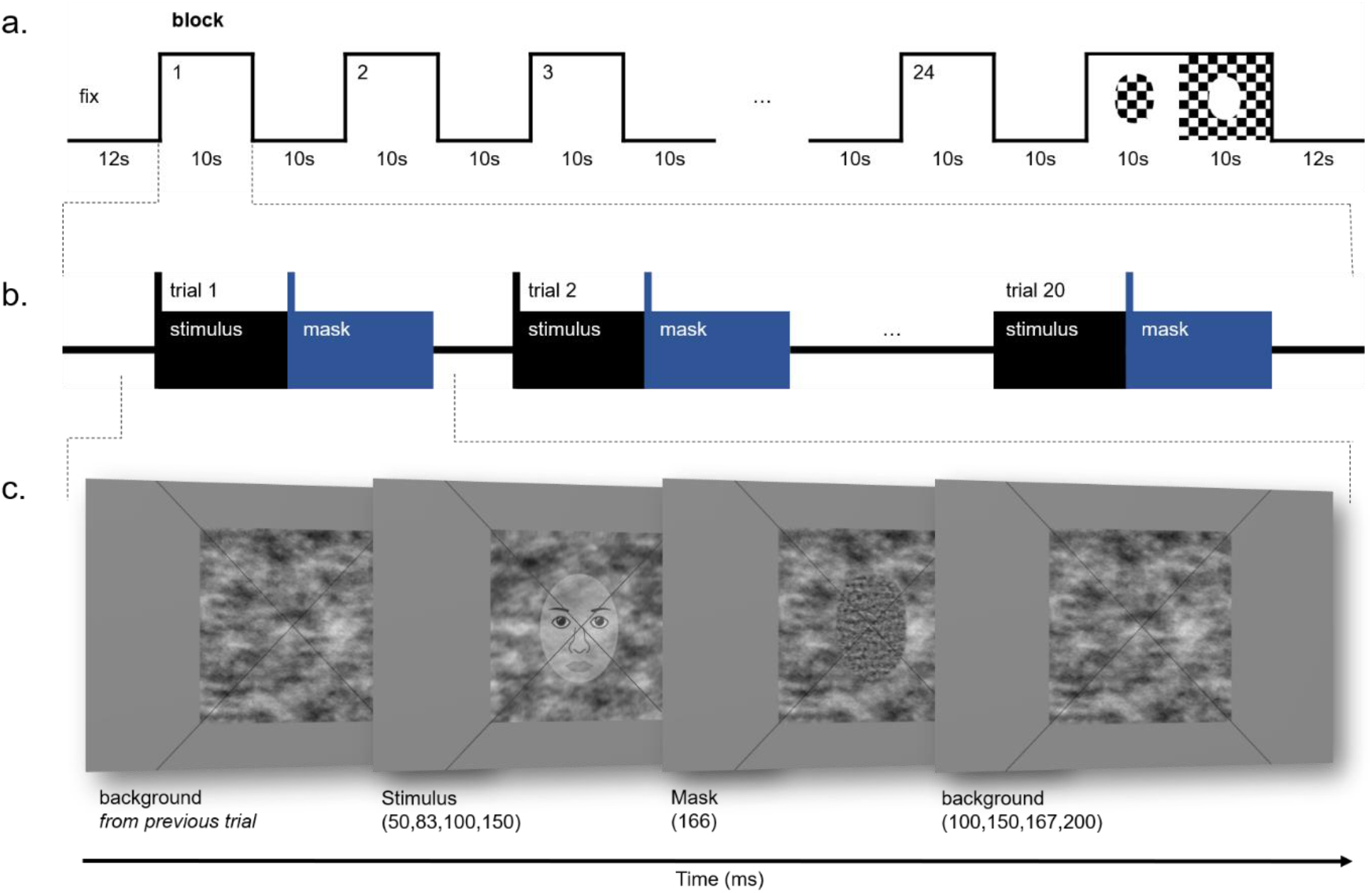
Design of main experiment. a) Every experimental run contained 24 blocks, alternated by fixation periods and ended with two retinotopic checkerboard blocks. b) A block consisted of 20 stimulus trials of the same experimental condition. c) An example trial: a face stimulus followed by a phase scrambled mask only containing HSFs. The alpha blending of the face stimulus was adapted for illustration purposes; experimental stimuli were less visible. Cartoon faces are used in replacement of photographs in the figure as journal regulations prohibit the use of identifiable faces.

Each block included 20 stimulus trials and four pseudo trials (417ms each; 25 frames at 16.67ms per frame) in which the background remained but no face or mask was presented. Two pseudo trials appeared at random during the first half of a block, and two more appeared at random during the second half. These were included to make the appearance of the stimuli less predictable. The other 20 trials contained stimuli from the same experimental condition. The order of the 20 stimulus trials in a block was randomised with the constraint that each trial was equally likely to appear at the beginning, middle and end of the block. The background remained stable within the block.

A stimulus trial consisted of a face stimulus presented for either 50, 83, 100 or 150ms, followed by a 166ms mask, after which only background remained (see Figure 2c). Each event duration within a trial was a multiple of the frame rate (16.67ms).

#### Functional localiser

During the 12.2-minute localiser run, subjects viewed greyscale images of either intact or scrambled faces, hands or instruments. Images from all categories were presented in 36 blocks in total (one image category per block), with six blocks per condition. A condition was never repeated twice in a row and the order of conditions was counterbalanced across subjects. A block comprised of 10 images in a random sequence, and each image was repeated three times throughout the functional localiser run. Each image appeared for 500ms, followed by a 500ms interstimulus interval.

### MRI acquisition

Subjects were scanned in a 3Tesla GE Signa Premier MRI scanner with a 48ch head coil, based at Cliniques Universitaires UCL Saint-Luc in Brussels. As anatomical references, whole-brain T1-weighted images were obtained during the first sessions (3D MP-RAGE, 1 × 1 × 1 mm, FOV = 256 mm, TI = 900ms, FA = 8°). Functional T2*-weighted GE echo-planar imaging was used to obtain the BOLD signal as an indirect measurement of neural activity. Thirty-two 2.4-mm axial slices were acquired (2.4mm isotropic, FOV = 240mm, TR = 2000ms, TE = 30ms, FA = 90°).

### Preprocessing

Functional and anatomical data were organised into the *Brain Imaging Data Structure* (Gorgolewski et al., 2016). Pre-processing of the data was carried out with *fMRIPrep 20*.*1*.*1* (Esteban et al., 2018; Esteban et al., 2019), which is based on *Nipype 1*.*5*.*0* (Gorgolewski et al., 2011; Gorgolewski et al., 2018). To ensure reproducibility using the specific software versions for *fMRIPrep* and all its dependencies, it was executed via its *Docker* container (Merkel, 2014).

#### Anatomical data pre-processing

Each T1-weighted (T1w) volume was corrected for intensity non-uniformity (Tustison et al., 2010; N4BiasFieldCorrection), and skull-stripping was executed (*antsBrainExtraction*.*shv*, OASIS30ANTs template). Next, brain tissue segmentation of cerebrospinal fluid (CSF), white-matter (WM) and grey-matter (GM) was performed on the brain-extracted T1w (using fast from FSL 5.0.9; Zhang et al., 2001). Finally, brain surfaces were reconstructed (using recon-all from *FreeSurfer* 6.0.1; Dale et al., 1999), and the brain mask estimated previously was refined with a custom variation of the method to reconcile ANTs-derived and *FreeSurfer*-derived segmentations of the cortical grey-matter of *Mindboggle* (Klein et al., 2017).

#### Functional data pre-processing

For each of the 24 functional runs per subject (across all tasks and sessions), the following pre-processing was performed. First, to generate a functional reference, volumes with substantial T1w contrast derived from nonsteady states of the scanner (volumes at the beginning of EPI sequence) were identified, realigned, and averaged. After skull-stripping of the functional reference volume, head motion parameters with respect to the functional reference (transformation matrices, and six corresponding rotation and translation parameters) were estimated (Jenkinson et al., 2002; *mcflirt* - FSL 5.0.9). On average, the maximum movement was 1.7 ± 0.32mm. After correcting for slice timing (Cox & Hyde, 1997; *3dTshift* from *AFNI 20160207*), the functional reference was co-registered to the T1w reference using boundary-based registration (*bbregister, FreeSurfer*; Greve & Fischl, 2009). The co-registration was configured with six degrees of freedom (i.e. 3 rotations and 3 translations).

Next, all functional data were resampled onto their original, native space by applying the transforms to correct for head-motion. Several confounding time series were calculated based on the functional data: framewise displacement (FD), DVARS and three region-wise global signals. FD was computed with the absolute sum of relative motions (*Power*, Power et al., 2014) and the relative root mean square displacement between affines (*Jenkinson*, Jenkinson et al., 2002). FD and DVARS were calculated for each functional run, both using their implementations in *Nipype* (Power et al., 2014). Frames that exceeded a threshold of 0.5 mm FD or 1.5 standardised DVARS were annotated as motion outliers. The three global signals were extracted within the CSF, the WM, and the whole-brain masks.

Using *FEAT* (FMRI Expert Analysis Tool; Version 6.0; FMRIB’s Software Library, www.fmrib.ox.ac.uk/fsl) the functional data were smoothed in the space domain using a Gaussian kernel of FWHM 5mm. And high-pass temporal filtering was carried out using a Gaussian-weighted least-squares straight-line fitting (sigma = 50.0s). For the whole-volume group-level analysis, we transformed for each subject the first-level estimate maps intact-scrambled for each condition into standard MNI152NLin2990c space with a resolution of 2mm3 using ANTs (*antsApplyTransforms*) employing the BSpline interpolation method. After the transformation, we smoothed the data with a gaussian kernel (FWHM = 8mm) using the *fslmaths* command.

### Regions of interest

V1 was delineated based on anatomical landmarks and functional data. We first localised V1 anatomically based on the *Freesurfer* atlas label (Figure 3c). Next, we selected the voxels showing a significantly stronger response to the stimulus region containing the face compared to its background. To do so, we contrasted the responses to the flickering checkerboard conditions in the block at the end of every run: [stimulus-background] (see Figure 3a). The voxels from this contrast (Figure 3b) that fell within V1, were defined as the V1 region of interest (ROI) in the current study (Figure 3d and Table 1).

**Table 1.** ROI MNI152 coordinates and volume sizes averaged over 16 subjects (with standard deviation). Coordinates are based on the median voxel in each ROI.

| *ROI* | *hemi* | *N voxels* | *x* | *y* | *z* |
| --- | --- | --- | --- | --- | --- |
| FFA | Right | 143(104) | 41(3.4) | -47(8.4) | -19(2.9) |
|  | Left | 108(93) | -27(2.8) | -52(9.4) | -14(3.1) |
| V1 | Right | 144(65) | 14(5.2) | -91(7.1) | -3(5.1) |
|  | Left | 155(53) | -11(6.0) | -93(2.4) | -8(4.1) |

**Figure 3.**
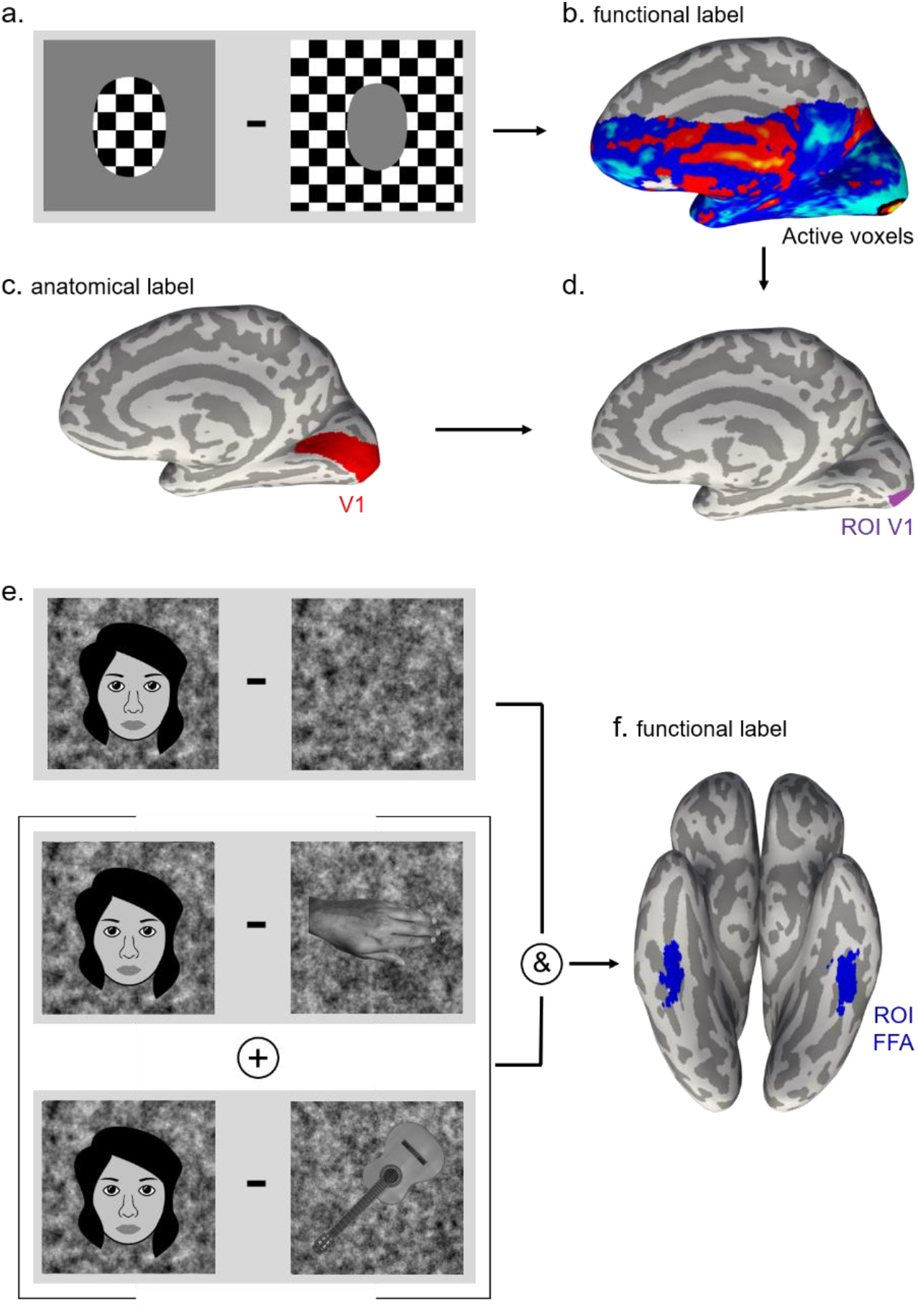
V1 and FFA ROI selection. a) Example checkerboard patterns of the retinotopy localiser. The pattern was confined to either the region where the face appeared in the main experiment or to the background. b) Contrasted [face checkerboards – background checkerboards] activation maps projected on an inflated brain surface of an example subject. A line is drawn to delineate the cluster of activated voxels in the (extra)striate cortex that show stronger activation due to the face checkerboards than the background checkerboards. c) The region of V1 was delineated using the Freesurfer atlas label. b) Inflated brain surface of an example subject with the final ROI consisting of voxels that fell both in the stimulated region (b) and the Freesurfer V1 region (c). e) Functional localisation stimuli examples. To rule out activity driven by the amplitude spectrum of face stimuli, we contrasted them with phase-scrambled stimuli. We then computed the conjunction of this cluster with a cluster more active to faces compared to hands or instruments. This way we ensured that we selected the brain region that is more receptive to faces than non-facial stimuli (Rossion et al., 2012). f) Example subject with the functionally defined FFA projected on the surface area of an inflated brain. Cartoon faces are used in replacement of photographs in the figure as journal regulations prohibit the use of identifiable faces.

The region responding preferentially to faces in the fusiform cortex (FFA) was identified independently for each subject, based on the functional localiser scan. First, to select the brain regions more responsive to faces than non-face objects we computed the conjunction between [intact faces - intact hands] and [intact faces - intact instruments] contrasts (Figure 3e). Next, to exclude activity merely driven by the amplitude spectrum of face images we selected the voxels that showed a significantly larger response to intact than scrambled faces [intact face - scrambled face] (Rossion et al., 2012). Significant voxel clusters on t maps were selected at a *q*[false discovery rate, FDR] < 0.01. After visual inspection, the threshold for some subjects was increased (cluster-threshold z-value > 3.0) to delineate between the fusiform and occipital face preferring clusters (see Figure 3f for an example subject). Since we were not able to delineate mid and posterior fusiform areas in all subjects, we did not distinguish between mFus- and pFus-faces in the current study (e.g. Weiner & Grill-Spector, 2013). Overall, and in line with past evidence (e.g. Kanwisher et al., 1997; Rossion et al., 2000), there were more face-selective voxels in the right FFA compared to the left (see Table 1). The voxel coordinates were in correspondence with previour research (see Table 1; Fahrenfort et al., 2012; Goffaux et al., 2011; Kanwisher et al., 1997; Rossion et al., 2003a; Thome et al., 2021). Additionally, we used the data obtained from the functional localiser to confirm that voxels in V1 showed greater responses to scrambled compared to intact faces (e.g. Goffaux et al., 2016). To test whether the [intact face - scrambled face] contrasted beta-values in V1 were significantly differing from 0, we conducted one-sample t-tests with a Bonferroni multiple comparison correction.

### Statistical Analyses

#### Behaviour analysis

In order to maintain attention to the stimulus throughout the experiment, subjects had to indicate whether the entire stimulus turned slightly red. Because of technical difficulties in the experimental code, the RT were not correctly recorded for all trials and could not be analysed.

We determined the accuracy for each condition per subject, ran a Linear Mixed effects Model (LMM) in R, and used a *Type III Analysis of Variance* (ANOVA; *Satterthwaite’s* approach) to determine whether there were differences in accuracy between conditions. We used a stepwise variable selection procedure with backward elimination (Heinze et al., 2018) and ended up with a model only including main effects (see Table 2).

**Table 2.** Overview of the LMMs after stepwise elimination of non-significant fixed effects. These final models were used to analyse either the accuracy of behavioural responses, ROI data (V1 or FFA), PPI data (between V1 and FFA) or individual voxels for the whole-volume analysis.

|  | *Model* | *BIC* |
| --- | --- | --- |
| Accuracy | Accuracy ~ 1 + SOA + SFmask + StimulusType + (1 \| subject) | -1443.545 |
| V1 | BetaValue ~ 1 + SOA * SFmask + SFmask * StimulusType + (1 \| subject) | 118624.6 |
| FFA | BetaValue ~ 1 + SOA * SFmask + SOA * StimulusType + (1 \| subject) | 107475 |
| PPI (V1-FFA) | BetaValue ~ 1 + StimulusType + (1 \| subject) | 28482.8 |
| Whole-volume | BetaValue ~ 1 + SOA * SFmask * StimulusType + (1 \| subject) |  |

#### ROI analysis

Individual beta weights were estimated for each condition of the SF experiment from each average ROI time course (left and right V1 and FFA). To separate the activity related to the processing of the face stimulus from general visual responses, we contrasted intact-scrambled in each experimental condition. These contrasted beta-values were used for the second-level analysis.

For each ROI in the second-level analysis (conducted in R), we used LMM approach to fit the contrasted beta-values from the first-level GLM, with fixed effects of SOA, SF masking, stimulus type, hemisphere and their interactions. Global variability between subjects was accounted for by modelling subject as a random effect (allowing individuals to have their own intercept about the group mean). For model fitting, we used the stepwise variable selection procedure, following backward elimination (Heinze et al., 2018). We started with the most complex model containing all variables and interaction terms (full model; outcome of the full models were used to plot the data for illustrative purpose), then we removed the non-significant four, three, and two-way interactions one by one to find the simplest model possible and thus maximising power in the analysis (for a detailed step-by-step overview of the fitted models, see Supplementary material 3). We validated the resulting model by determining whether it yields the smallest Bayesian information criterion (BIC).

In both ROIs, there was no significant effect of hemisphere, so we collapsed the data from the two hemispheres. The final model for V1 included a two-way interaction term of SF masking and stimulus type, as well as SF masking and SOA (Table 2). The final model for FFA included two-way interaction terms of SOA with stimulus type, and SF masking with SOA (Table 2).

To interpret the estimated means we used the “*stats*” package in R to run a Type III ANOVA using Satterthwaite’s method for degrees of freedom. Since we expected to see the cross-like pattern of differential activity, i.e. high differential responses for the early disruption of LSF, decreasing as a function of SOA and the opposite for HSF masking, we obtained slope values for the different conditions (with the *emtrends* function of the *emmeans* package). The difference in the slope values was then tested using a pairwise comparison approach (*pairs*).

#### Whole-volume group-level analysis

To identify other regions involved in coarse-to-fine processing, we performed a whole-volume group-level analysis, again using a LMM approach to fit the contrasted beta-values from the first-level GLM (i.e., outcome of intact-scrambled contrast). Here, we fit the same model on every voxel in the fMRI data and, unlike the models fit for the ROIs, retained all interaction terms regardless of whether or not they were significant. This is because different voxels would have required different terms to be removed and thus reduced the interpretability of the resulting statistical map (since each voxel would have been modelled differently). Therefore, we opted in favour of interpretability of the results (possibly sacrificing power for lower order interaction terms in models with redundant higher order interaction terms). For every voxel, we fit the full LMM (using *R*) including all interaction effects (Table 2).

We performed an ANOVA on each voxel’s model predictions to obtain a p-value for each term, resulting in p-value maps. We adjusted the p-value maps for multiple comparisons using FDR correction (in *R*, using the *p*.*fdr* function from the *FDRestimation* package with adjust.method=“BH”). To ensure that the voxels with an SOA by SF masking interaction effect indeed exhibit the ‘coarse-to-fine signature’ (i.e. more positive slopes when accumulating HSF than LSF), we obtained the slope values for the LSF and HSF masking conditions as a function of SOA. We subtracted the LSF from the HSF slopes, such that positive values support the ‘coarse-to-fine signature’. There were no voxels with a negative slope difference (i.e., no significant ‘fine-to-coarse signature’). To visualise the results on the cortical surface, we projected the difference map of the fitted slopes for LSF and HSF and thresholded this map with the FDR corrected p-value map.

#### Psychophysiological Interaction

To address whether the activity between V1 and FFA is coupled more strongly when coarse-to-fine processing can take place compared to when we disrupt it via masking, we conducted a psychophysiological interaction (PPI) analysis. The average time-series of the V1 ROI was used as the seed and was extracted using *fslmeants* to form a physiological regressor. A psychological regressor was created for each condition by convolving a double gamma HRF with the task regressor (O’Reilly et al., 2012). The PPI regressors were created by multiplying the physiological and psychological regressors (one per condition). The model also incorporated the six motion regressors. For all subjects, the different regressors were entered into a GLM, and the PPI interaction terms were estimated in the FFA.

Next, we applied the LMM stepwise variable selection procedure with backward elimination, similar to the main analysis. Starting with the most complex model, and removing non-significant interaction terms to yield the simplest model that accounted for the data (Table 2).

## Results

### Behavioural results

Subjects performed an easy colour change detection task, which randomly occurred twice per block. The performance accuracy was 97.02% (SD = 0.009). After fitting an LMM with full interaction effects and pruning this model to the simplest model with only main effects, the ANOVA showed that there were no significant differences of accuracy between any of the conditions (p-values ranging between .234 and .588; Table 3). The global variance between subjects only explained a small proportion of the residual variability (0.1%).

**Table 3.** Type III ANOVA - for accuracy of behavioural responses and ROI data in V1 and FFA.

|  | *Sum Sq* | *Mean Sq* | *NumDF* | *DenDF* | *F* | *p* |  |
| --- | --- | --- | --- | --- | --- | --- | --- |
| <u>Accuracy</u> |  |  |  |  |  |  |  |
| SOA | 0.00000005 | 0.00000005 | 1 | 364 | 0.0000 | 0.9963 |  |
| SFmask | 0.00013665 | 0.00013665 | 1 | 364 | 0.0648 | 0.7992 |  |
| StimulusType | 0.00196569 | 0.00098284 | 2 | 364 | 0.4660 | 0.6279 |  |
| <u>V1</u> |  |  |  |  |  |  |  |
| SOA | 3110 | 3110 | 1 | 10187 | 0.4808 | 0.489 |  |
| SFmask | 154801 | 154801 | 1 | 10187 | 23.9336 | <.001 | *** |
| StimulusType | 7911 | 7911 | 1 | 10187 | 1.2231 | 0.269 |  |
| SOA:Sfmask | 192966 | 192966 | 1 | 10187 | 29.8342 | <.001 | *** |
| SFmask:StimulusType | 21943 | 21943 | 1 | 10187 | 3.3926 | 0.066 | . |
| <u>FFA</u> |  |  |  |  |  |  |  |
| SOA | 1273009 | 1273009 | 1 | 10187 | 589.4631 | <.001 | *** |
| SFmask | 52022 | 52022 | 1 | 10187 | 24.0887 | <.001 | *** |
| StimulusType | 2 | 2 | 1 | 10187 | 0.0009 | 0.976 |  |
| SOA:Sfmask | 58632 | 58632 | 1 | 10187 | 27.1492 | <.001 | *** |
| SOA:StimulusType | 66649 | 66649 | 1 | 10187 | 30.8614 | <.001 | *** |

### ROI results

#### V1

First, to verify that the intact-scrambled response difference was negative in less constrained (i.e., unmasked and less rapid) stimulation conditions, we compared responses of intact and phase-scrambled faces from the functional localiser in V1. We found that the intact-scrambled response difference (M = -.16, SD = .26) was significantly lower than zero (t(31) = -3.56, p = 0.002).

In the main experiment, we predicted that the early masking of LSF would result in a quasi null intact-scrambled difference, in line with the hypothesis that early LSF processing guides the formation of a sharp representation in V1, and preventing such guidance results in a heightened response to intact faces. However, at 50 ms of SOA, the intact-scrambled difference was positive (see Figure 4) indicating that the response in V1 was larger for intact than for scrambled faces. Although this may appear puzzling, it could reflect the stimulus heterogeneity of intact blocks, and thus stronger stimulus on- and offset responses than in scrambled blocks. LSF masking caused progressively less disturbance over time as shown by the progressive decrease of the intact-scrambled differential response. At 150 ms, the response differential reached negative values, indexing the successful formation of a face representation in V1. The reverse pattern was observed when masking HSF, creating the hypothesized cross-like pattern of SF masking, the expected ‘signature’ of coarse-to-fine processing in V1. These observations were borne out by the LMM (Figure 4).

**Figure 4.**
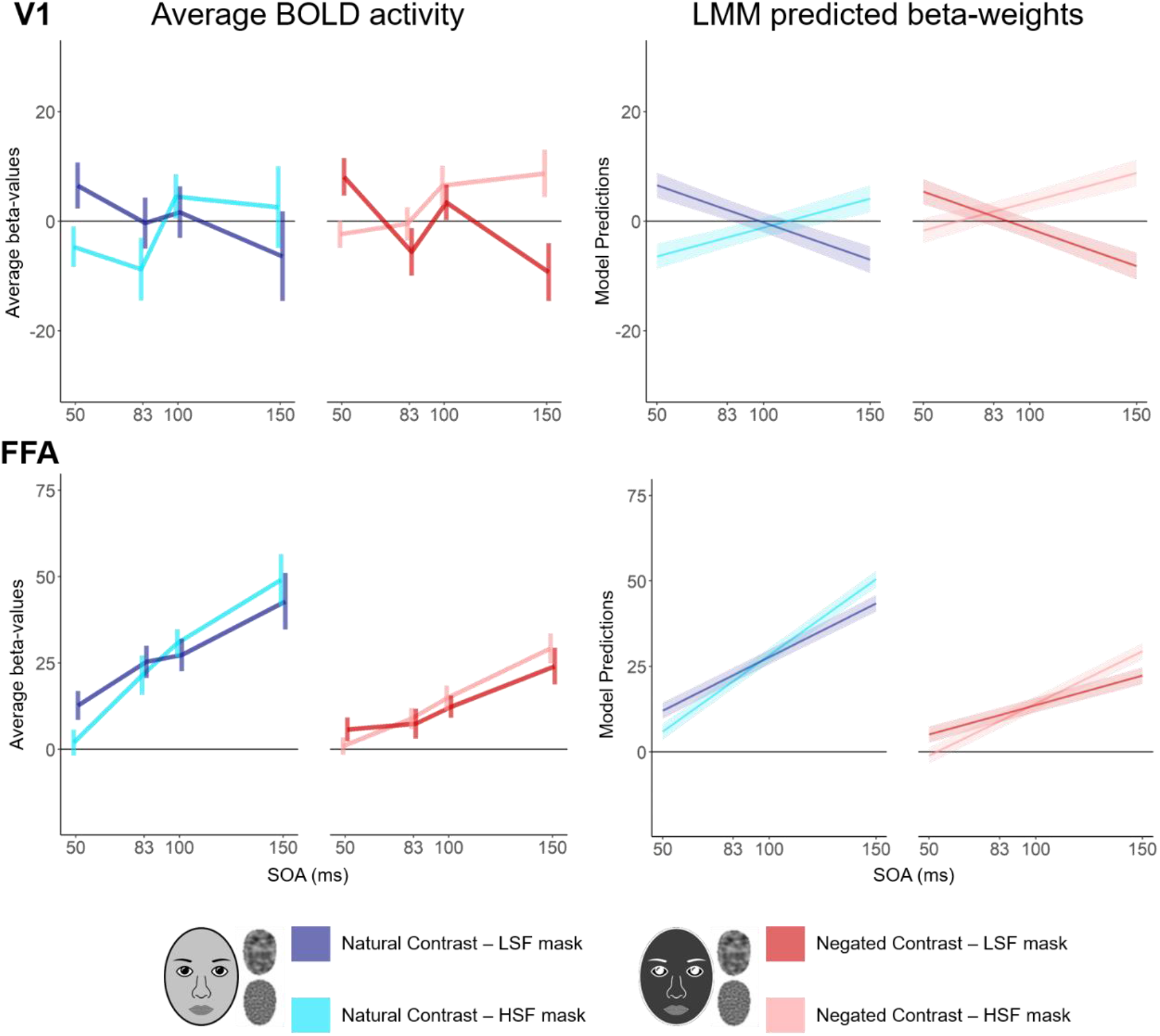
Intact-scrambled response differential for both V1 (top) and FFA (bottom). Plots in the left column represent the group averaged beta-values including a 95% within-subjects confidence interval as described in Loftus & Masson, 1994. The LMM predictions shown on the right plots are from a full model including all interaction effects which was ran just for illustrative purposes. In both V1 and FFA, there was a significant interaction effect of SOA with SF masking. The FFA showed an additional interaction effect of SOA with stimulus type. Cartoon faces are used in replacement of photographs in the figure as journal regulations prohibit the use of identifiable faces.

After systematically removing non-significant higher-order interaction terms (as determined by a *type III ANOVA*), the LMM included two-way interaction effect terms of SF masking with SOA and of SF masking with stimulus type (see Table 3). Only a negligible proportion of the residual variability was accounted for by the global differences between subjects (0.001%). Table 3 summarises the significant terms in the LMM. Since we did not find any hemispheric differences, we collapsed the data over both hemispheres in the LMM for more power in the analysis. We found a significant interaction effect between SOA and SF masking, and a marginally significant interaction effect between SF masking and stimulus type, the latter we did not further analyse since it was not statistically significant. Figure 4 depicts the average BOLD activity and the LMM predictions from the full model, although not all terms were significant we include them in the figure for illustrative purposes.

For both naturally contrasted and negated images, we found that in V1 the intact-scrambled response differential depicted a cross-like pattern. There was a negative slope as a function of SOA when masking LSFs (collapsed over stimulus type: β = -.14, CI = .12), whereas HSF masking yielded a positive slope (β = .11, CI = .12). These slopes differed significantly from each other (z = -5.46, p < .001).

Table 4 shows the estimated marginal means of the model as well as pairwise comparisons between LSF and HSF masked conditions. V1 response differential was significantly bigger when disrupting LSFs compared to HSFs at 50ms. By 150ms, this trend was reversed with disruption of HSFs causing a significantly larger V1 response than disruption of LSFs (see Table 4).

**Table 4.** Model predictions (estimated marginal means - emmeans) and pairwise comparisons on predicted variables between LSF and HSF masked conditions for different SOAs in both V1 and FFA.

|  |  | *LSF mask* | *HSF mask* |  |  |  |
| --- | --- | --- | --- | --- | --- | --- |
| *ROI* | *SOA* | *emmean (SE)* | *emmean (SE)* | *Estimate (SE)* | *Test statistic (z/t)* | *p* |
| V1 |  |  |  |  |  |  |
|  |  |  | <u>Natural</u> |  |  |  |
|  | 50 | 6.56 (2.28) | -6.40 (2.28) | 13.00 (3.02) | 4.29 | <.001 |
|  | 83 | 2.08 (1.82) | -2.93 (1.82) | 5.02 (2.32) | 2.16 | 0.031 |
|  | 100 | -0.22 (1.79) | -1.15 (1.79) | 0.92 (2.26) | 0.41 | 0.683 |
|  | 150 | -7.01 (2.46) | 4.11 (2.46) | -11.10 (3.28) | -3.38 | <.001 |
|  |  |  | <u>Negated</u> |  |  |  |
|  | 50 | 5.39 (2.28) | -1.71 (2.28) | 7.10 (3.02) | 2.35 | 0.019 |
|  | 83 | 0.91 (1.82) | 1.76 (1.82) | -0.85 (2.32) | -0.37 | 0.715 |
|  | 100 | -1.40 (1.79) | 3.55 (1.79) | -4.94 (2.26) | -2.19 | 0.029 |
|  | 150 | -8.18 (2.46) | 8.80 (2.46) | -17.00 (3.28) | -5.17 | <.001 |
| FFA |  |  |  |  |  |  |
|  |  |  | <u>Natural</u> |  |  |  |
|  | 50 | 12.11 (2.36) | 5.95 (2.36) | 6.16 (1.48) | 4.15 | <.001 |
|  | 83 | 22.46 (2.15) | 20.68 (2.15) | 1.78 (0.98) | 1.82 | 0.068 |
|  | 100 | 27.79 (2.13) | 28.27 (2.13) | -0.48 (0.93) | -0.517 | 0.605 |
|  | 150 | 43.47 (2.44) | 50.59 (2.44) | -7.11 (1.66) | -4.29 | <.001 |
|  |  |  | <u>Negated</u> |  |  |  |
|  | 50 | 5.11 (2.36) | -1.05 (2.36) | 6.16 (1.48) | 4.15 | <.001 |
|  | 83 | 10.79 (2.15) | 9.01 (2.15) | 1.78 (0.98) | 1.82 | 0.068 |
|  | 100 | 13.72 (2.13) | 14.2 (2.13) | -0.48 (0.93) | -0.517 | 0.605 |
|  | 150 | 22.32 (2.44) | 29.44 (2.44) | -7.11 (1.66) | -4.29 | <.001 |

#### FFA

As with V1, the proportion of residual variance explained by the random effect of subject was negligible (0.028%) and there was no interaction nor main effect of hemisphere; we therefore collapsed the data over left and right FFA. The LMM in FFA included the interaction effect between SOA and stimulus type, as well as between SOA and SF masking.

Unlike V1, the intact-scrambled response differential was positive in the FFA as this region is known to respond more to intact faces than scrambled faces (e.g. Rossion et al., 2012). Irrespective of the masked SF, the differential intact-scrambled response increased as a function of SOA. The influence of SOA was stronger for naturally contrasted faces (β = .38, CI = .07) than for contrast negated faces (β = .24, CI = .07). The slopes differed significantly (z = -5.56, p < .001), and there was a significant interaction between SOA and stimulus type (see Table 3 and Figure 4).

We observed the hypothesised coarse-to-fine signature in the form of a significant interaction between SOA and SF masking; similar to V1, the fitted slope was more positive when masking HSF (β = .38, CI = .07) than LSF (β = .21, CI = .07; test of the slope difference: z = -5.21, p < .001).

### Whole-volume group-level results

In a whole-volume group-level LMM analysis, we investigated potential coarse-to-fine dynamics outside of V1 and FFA. The resulting p-value maps were FDR corrected for multiple comparisons. The only 2-way interaction term that showed significance in the whole-volume analysis was that of SOA and SF masking and there were no voxels with significant 3-way interactions. Several regions showed this interaction (see Figure 5 and Table 5). Along the ventral visual stream, the interaction was found significant in the right fusiform gyrus (BA 37), confirming findings from the ROI analysis. We did not find the effect to be significant in the left fusiform gyrus. The SOA by SF masking interaction was also significant in bilateral parietal and ventrolateral/orbitofrontal clusters (e.g., middle temporal gyrus or BA 21; inferior temporal gyrus or BA 20; see Table 5) - which have been theorised to be an origin of the coarse template-based feedback in coarse-to-fine processing (e.g., Bar, 2004; Bullier, 2001). In all cases, these 2-way interactions were due to a pattern reminiscent of what we already observed in the ROI analyses for V1 and FFA (Figure 4): a more positive slope with the progressive accumulation of HSF compared to LSF (HSF-masking slope: β = .062, CI = .002; LSF-masking slope: β = -.089, CI= .002; test of the slope difference: z = -197.71, p < .001). Specifically, at 50ms the intact-scrambled response differential was larger when disrupting the LSFs compared to HSFs, with the reverse being true at 150ms, in line with the anticipated coarse-to-fine signature. The effect of stimulus type (faces with natural vs negated contrast) did not survive the multiple comparisons correction (Fs < 9.35, q[FDR] > .95).

**Table 5.** Whole-volume group-level analysis. Regions showing the interaction effect of SF masking with SOA (CtF signature). For each cluster, the hemisphere (R = right hemisphere; L = left hemisphere), Brodmann’s area (BA), number of voxels (k), and MNI152 coordinates (x, y, z). Based on the median voxel in each cluster of voxels.

| *Cluster nr* |  | *hemi* | *BA* | *N voxels* | *x* | *y* | *z* |
| --- | --- | --- | --- | --- | --- | --- | --- |
| Frontal areas |  |  |  |  |  |  |  |
| 4 | Frontal_Inf_Orb | R | 47 | 131 | 29 | 35 | -8 |
| 6 | Frontal_Inf_Orb | L | 47 | 1152 | -35 | 37 | -10 |
| 20 | Frontal_Inf_Tri | L | 46 | 20 | -35 | 34 | 13 |
| 7 | Frontal_Med_Orb | L | 10 | 301 | -9 | 54 | -7 |
| 12 | Frontal_Sup_Orb | L | 10 | 134 | -16 | 57 | -8 |
| 2 | Postcentral | R | 4 | 145 | 48 | -6 | 27 |
| 23 | SupraMarginal | R | 40 | 4 | 51 | -23 | 36 |
| 26 | Postcentral | L | 4 | 45 | -48 | -14 | 32 |
| Temporal areas |  |  |  |  |  |  |  |
| 5 | Fusiform | R | 37 | 650 | 40 | -52 | -15 |
| 7 |  |  | 36 | 396 | 35 | -30 | -17 |
| 11 |  |  | 20 | 315 | 41 | -15 | -23 |
| 28 | Fusiform | L | 19 | 2 | -26 | -75 | -3 |
| 14 | Temporal_Mid | R | 21 | 104 | 45 | -26 | -7 |
| 4 | Temporal_Mid | L | 21 | 856 | -57 | -38 | -9 |
| 19 |  |  | 21 | 111 | -59 | -13 | -21 |
| 9 | Temporal_Mid | R | 39 | 263 | 40 | -57 | 16 |
| 20 | Temporal_Inf | R | 21 | 14 | 51 | -12 | -26 |
| 14 | Temporal_Inf | L | 20 | 203 | -45 | -25 | -21 |
| 21 |  |  | 38 | 13 | -42 | 3 | -33 |
| 3 | Temporal_Pole_Mid | L | 38 | 410 | -35 | 11 | -31 |
| Occipital areas |  |  |  |  |  |  |  |
| 1 | Lingual | R | 19 | 2012 | 18 | -91 | -5 |
| 8 | Occipital_Mid | R | 39 | 85 | 29 | -69 | 23 |
| 10 |  |  | 18 | 169 | 29 | -83 | 8 |
| 23 | Occipital_Mid | L | 19 | 28 | -29 | -72 | 20 |
| 13 |  |  | 19 | 74 | -33 | -81 | 18 |
| 8 |  |  | 39 | 54 | -35 | -75 | 38 |
| 16 | Occipital_Sup | R | 47 | 77 | 29 | -60 | 31 |
| 17 |  |  | 19 | 84 | 23 | -83 | 19 |
| 21 |  |  | 18 | 12 | 22 | -94 | 16 |
| 5 | Occipital_Inf | L | 18 | 1875 | -20 | -95 | -4 |
| 15 | Calcarine | R | 17 | 25 | 9 | -91 | 11 |
| 1 | Calcarine | L | 17 | 259 | -9 | -88 | 10 |
| 27 | Lingual | L | 19 | 12 | -15 | -51 | -2 |
| 15 |  |  | 19 | 13 | -25 | -72 | -2 |
| Parietal areas |  |  |  |  |  |  |  |
| 0 | Parietal_Inf | L | 39 | 3466 | -40 | -49 | 40 |
| 3 | Insula | R | 13 | 1636 | 35 | -8 | 17 |
| 16 | Insula | L | 13 | 75 | -33 | 0 | 12 |
| 25 |  |  | 13 | 44 | -34 | 7 | -11 |
| 9 |  |  | 1 | 286 | -34 | -24 | 22 |
| 19 | Angular | R | 39 | 10 | 41 | -58 | 44 |
| 12 |  |  | 39 | 174 | 37 | -67 | 43 |
| 17 | Angular | L | 39 | 42 | -39 | -64 | 38 |
| 29 | Precuneus | L | 31 | 6 | -17 | -41 | 47 |

**Figure 5.**
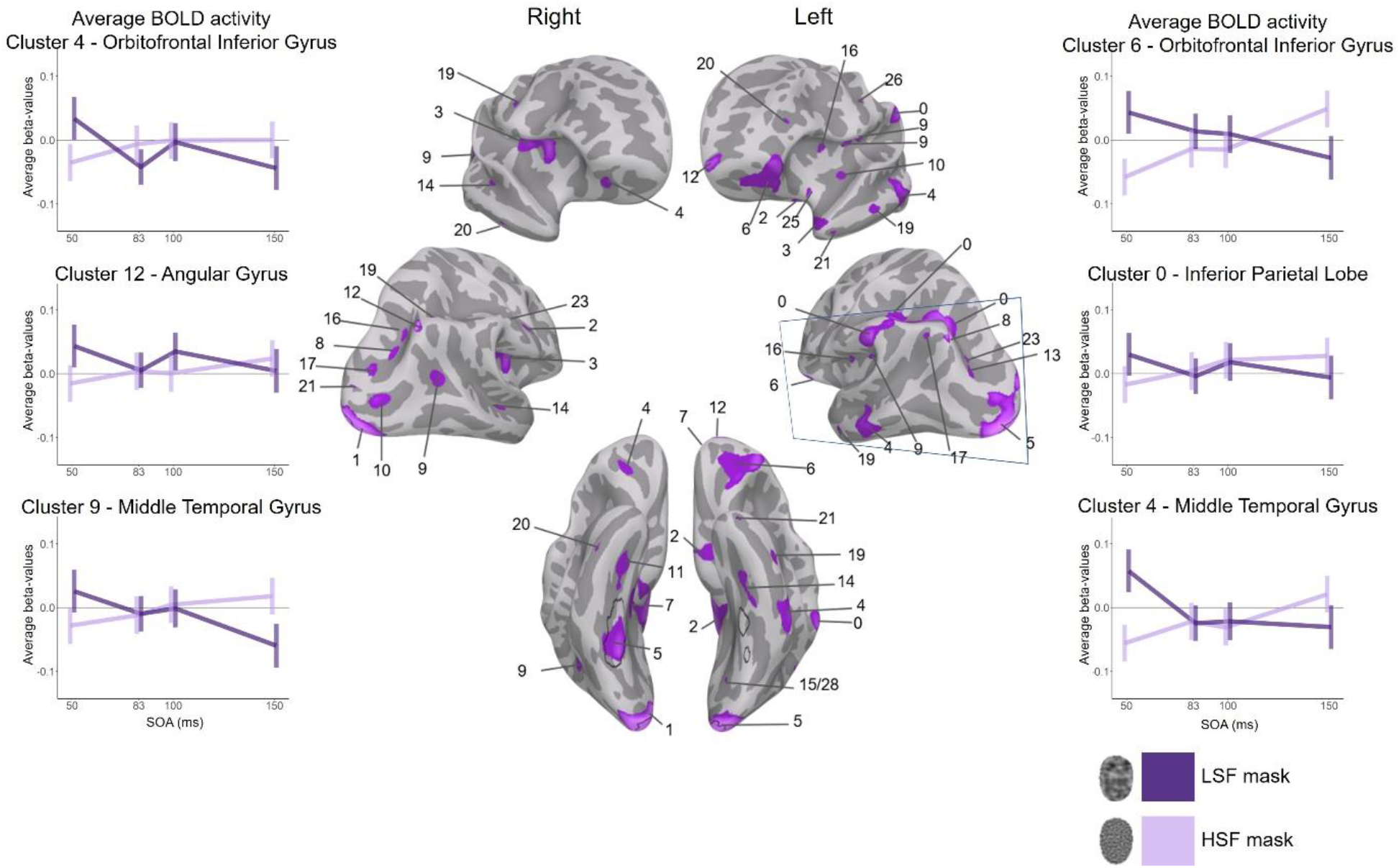
Whole-volume group-level analysis interaction map projected on an inflated brain surface. The thin box represents the brain volume that was scanned. Here we show the voxels with a significant SOA by SF masking interaction effect in the scanned volume. Purple highlighted areas show the difference in the fitted slope (HSF minus LSF) such that positive values support the CtF signature (LSF masking more disruptive when presented early than late = more negative slope and vice versa for HSF masking). The map was FDR corrected for multiple comparisons, and no surviving vertices showed a positive value for the slope difference (i.e. we did not observe any significant ‘fine-to-coarse signature’). For comparison with our ROI analyses, the black outlines indicate the location of the functionally defined V1 and FFA - collapsed over all subjects. The averaged beta-values in the plots are normalised.

### Psychophysiological Interaction

A premise of coarse-to-fine theories is that visual perception involves recurrent interactions between V1 and higher level regions in dorsal, frontal, and ventral brain regions. FFA has been shown to be particularly sensitive to any disruption of the natural statistics of the upright human face (e.g., Schiltz & Rossion, 2006; Yue et al., 2013), suggesting it contains a template of the human face presumably guiding coarse-to-fine integration. We thus tracked the influence of SF masking on the PPI connectivity between V1 and FFA.

The final LMM after pruning, however, only included stimulus type as fixed effect. There was just a small proportion of residual variability accounted for by the global differences between subjects (.004%). We found no effect of SOA nor of SF masking. The connectivity between V1 and FFA was stronger for naturally contrasted faces than for negated faces (F = 33.69, p < .001, see Figure 6). This effect cannot be attributed to differences in response amplitude between natural and negated faces, and suggests that the low and high tiers of the ventral pathway cooperate more strongly when the stimulus matches the internal template representation (i.e., face in a natural contrast polarity).

**Figure 6.**
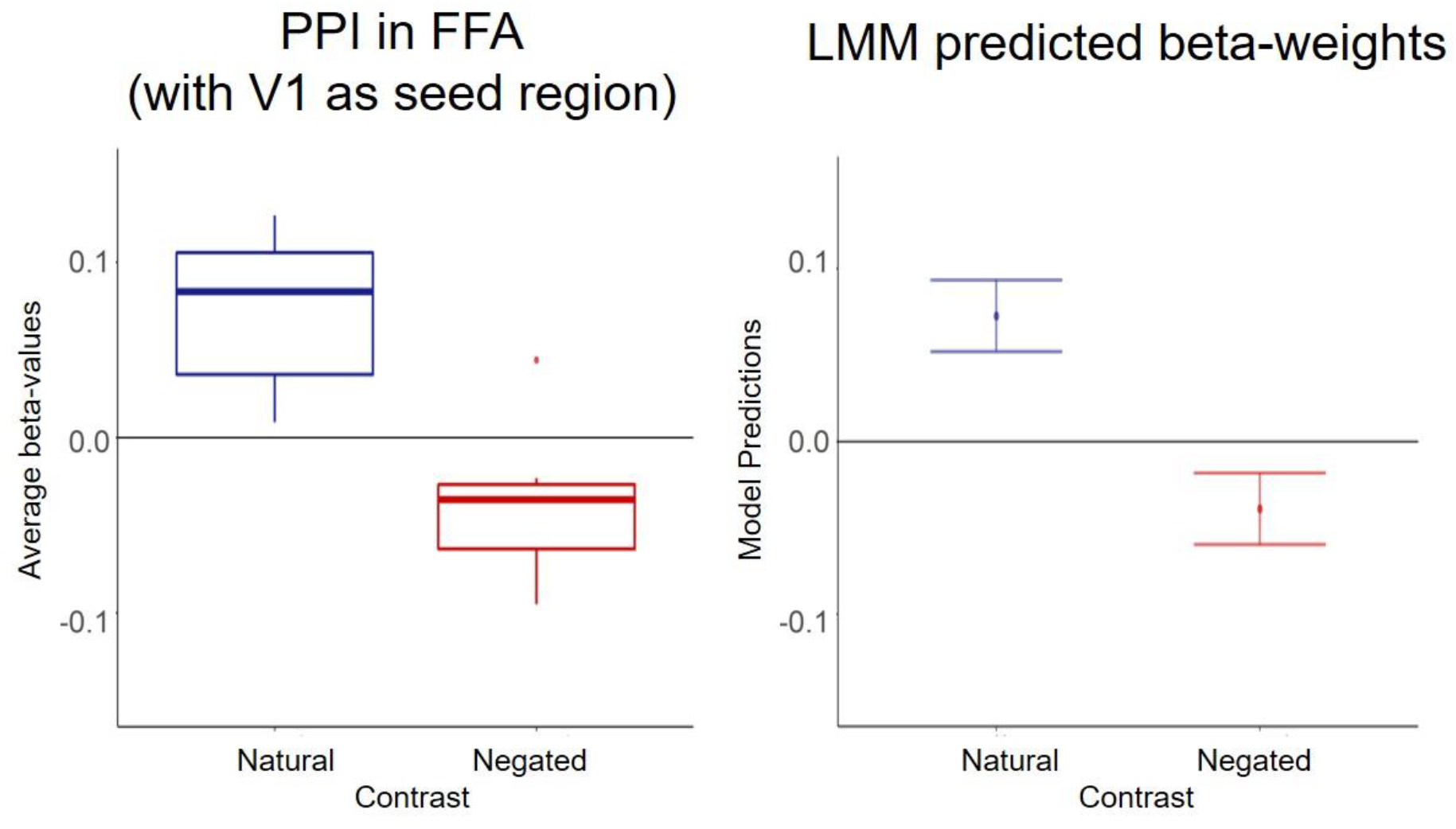
Results of the PPI analysis in the FFA with V1 as seed region. Left. Boxplots of the main effect of stimulus type on the PPI connectivity: upper and lower hinges correspond to the first and third quartile, winkers represent a 95% within-subject confidence interval (Loftus & Masson, 1994) and the single dot represents an outlier. Right. Boxplots showing the model predictions of the main effect of stimulus type (with 95% confidence interval) of a model with only stimulus type as main effect.

## Discussion

The notion that recurrent processing contributes to the refinement of the coarse representations initially extracted by the visual system (Kok & de Lange, 2014) is influential in vision and computer sciences (Bar, 2004; Bullier, 2001; Kreiman & Serre, 2020; Mohsenzadeh et al., 2018). In this framework, it has been proposed that V1 plays the role of an ‘active blackboard’ for the coarse-to-fine integration of visual input (Bullier, 2001; Deco & Lee, 2004; Roland, 2010). We reasoned that if coarse-to-fine processing is implemented in V1 as an efficient way to represent visual information, its disruption should have measurable costs, and result in suboptimal representations in this region. We disrupted coarse-to-fine integration via backward SF masking to prevent the contribution of coarse and fine spatial scales over the course of human face processing. Disrupting LSF processing early was predicted to disrupt top-down LSF-driven signals, prevent the formation of a sparse representation, and therefore increase V1’s response to the face stimulus. Our results support this prediction. Since LSFs contribute less to processing over time, disrupting LSF late was expected to be less disruptive, which was also supported by our results. The opposite trend was observed for HSF masking in line with the notion that fine cues, i.e. HSFs, are processed at a later stage, contributing progressively more to the sharpening of visual representations. In line with this notion, we found that the interference caused by HSF masking on responses in V1 increased over time. Our finding that the V1 response to strictly identical sets of full-spectrum face images follows opposite trends depending on the masked SF is compelling evidence of the complementary contribution of the coarse and fine scales to the build-up of progressively finer visual representations in V1 (e.g., Bar, 2004; Bullier, 2001; Hegde, 2008). By retaining a core aspect of naturalistic vision; i.e. full range of aligned SF, our work goes beyond past evidence of a coarse-to-fine precedence, and yields novel insights on a fundamental tenet of coarse-to-fine models, namely that the integration of visual information operates in a coarse-to-fine manner in V1.

Despite fMRI having a temporal resolution of several seconds, past and present results confirm that this technique can be used to address time-sensitive questions if combined with time-sensitive methods such as masking (e.g. Goffaux et al., 2011; Green et al., 2005; Grill-Spector et al., 2000; Haynes et al., 2005). The backward masking of specific SF ranges allowed us to address when and how the coarse and fine visual cues contribute to the build-up of face representations in V1. The drawback of our approach is that it is limited to a few *a priori* SOAs in contrast to other techniques like EEG and MEG that allow for a denser temporal sampling. Hansen et al. (2021) recently developed an EEG encoding-based mapping technique to map the variance of human visual ERPS onto the variance in location-specific SF content of a large dataset of natural scenes. They observed a coarse-to-fine shift in visual encoding over the first ∼150ms, which validates the time window used in the present experiment. The authors reported additional coarse-to-fine coding phases starting ∼180ms and 260ms after stimulus onset, suggesting that longer SOAs should be used in future studies.

The top-down signal driving the progressive sharpening of V1 representations is presumably a coarse representation of the visual input stored in more anterior high-level visual regions, the large receptive fields of which enable a more global representation of the stimulus (e.g., Lee, 2015). This tenet is supported by past findings that the high-level ventral regions found to preferentially respond to faces or places attune to LSFs more rapidly than to HSF content of their preferred stimulus class (Goffaux et al., 2011; Kauffmann et al., 2015c; Musel et al., 2014; Peyrin et al., 2006; Schyns & Oliva, 1994). Here, we show that, in full-spectrum viewing conditions, the disruption of coarse-to-fine integration modulates face processing in the human FFA similarly to V1. Indeed, despite the fact that the FFA diverges from V1 in its overall bigger response to intact faces than to scrambled faces and its steady increase with SOA (i.e., face visibility; e.g., Kay & Yeatman, 2016; Yue et al., 2013), the response in FFA increases more rapidly upon accumulation of meaningful HSF than LSF content, as has been found in the primary visual cortex. The whole-volume results confirmed that, in the ventral pathway, coarse-to-fine integration mainly involves V1 and FFA (for the stimuli used in our experiment). However, when localized at the individual level, we also found the OFA temporal profile to mirror the V1 coarse-to-fine dynamics (see Supplementary material 5). Considering the closer proximity to V1 and the existence of denser connections between V1 and OFA than FFA (Finzi et al., 2021), it may come as a surprise that this region did not show up in the whole-volume analysis. Consistent evidence has nonetheless indicated that the OFA encodes a finer scale of face information than FFA (e.g. Goffaux et al., 2011; Goffaux et al., 2012; Rossion et al., 2003a; Schiltz & Rossion, 2006), suggesting that, in contrast to FFA who would involve from the earliest LSF-driven stages, OFA may mainly contribute to the later stages of face representation refinement (e.g., Gentile et al., 2017; Rossion et al., 2003a).

We used contrast negation to address whether the observed coarse-to-fine dynamics involve a priori knowledge of the natural contrast polarity of the human face (e.g., Bar, 2004; Mumford, 1992). We replicate past findings that negation substantially hampers the accumulation of sensory input necessary at high-level stages of processing (Liu-Shuang et al., 2022; see also Bex & Makous, 2002). However, the coarse-to-fine dynamics observed in V1 (and FFA) were not influenced by negation. This suggests that the (coarse-to-fine) sharpening of face representation in V1 is guided by high-level neural populations that are sensitive to phase-aligned content irrespective of whether the input matches the natural contrast statistics of an internally stored category. Still, the significant PPI connectivity between V1 and FFA for naturally contrasted faces indicates that these regions interact during the formation of face-specialized representations. Considering V1’s relative immunity to negation (George et al., 1999; Yue et al., 2013), the modulation of the V1/FFA connectivity by negation is likely to result from a reduced top-down influence of FFA on primary visual processing when the stimulus fails to correspond to the internal template representation of a face. Yet, the fact that negation did not influence the coarse-to-fine dynamics in V1 suggests these are driven by other regions than the FFA, as supported by the whole-volume results (see below). Future investigation is however needed to support this interpretation of the data since our study may simply be underpowered to reveal the influence of contrast negation on the coarse-to-fine dynamics of V1 and on its connectivity to FFA.

Besides the FFA, our whole-volume analyses showed a coarse-to-fine signature in orbitofrontal and dorsal (parietal) regions. This agrees with growing evidence that object recognition crucially relies on the involvement of dorsal and frontal regions (Ayzenberg & Behrmann, 2022; Bar, 2004; Kay & Yeatman, 2016; Zachariou et al., 2017). These regions are thought to rapidly generate a global and coarse representation of incoming input, i.e. an initial ‘guess’ (Ayzenberg & Behrmann, 2022; Ayzenberg et al., 2022; Bar et al., 2006; Fintzi & Mahon, 2014; Kauffmann et al., 2015a; Kauffmann et al., 2015c; Kveraga et al., 2007; Peyrin et al., 2010; Zachariou et al., 2017). This is supported by past findings that the dorsal pathway represents shape before - and sends information to - the ventral pathway (Ayzenberg & Behrmann, 2022; Ayzenberg et al., 2022; Van Dromme et al., 2016). During human face processing, in particular, the posterior parietal cortex has been proposed to represent the global configuration of facial features and convey it to the face-selective ventral regions (Zachariou et al., 2017). In line with the absence of a negation effect upon V1’s coarse-to-fine dynamics, our results suggest that the coarse structure guiding the representation sharpening in V1 mainly comes from dorsal and orbitofrontal non face-selective populations. Altogether our findings therefore challenge the hypothesis that coarse-to-fine integration occurs primarily in inferotemporal cortex (e.g., Bar, 2004; Kveraga et al., 2007; Wyatte et al., 2014). They instead favour the notion that the coarse representation activated early in dorsal and orbitofrontal cortices is integrated with the representation of the fine details in V1. In this context, the inferotemporal cortex may not play an active role in coarse-to-fine integration but would encode the output signals delivered by the dorso-frontal-V1 coarse-to-fine network (see Deco & Lee, 2004 for a similar proposal).

The finding that masking of specific SFs differentially impacts V1 over processing course indicates that the contribution of this region to visual processing is sustained, which challenges the still influential assumption that V1’s role is limited to an early filtering of visual information to downstream regions (Kravitz et al., 2013). Our findings favour the proposal that V1 integrates incoming input with intrinsic and top-down orbitofrontal and dorsal influences. They add to the body of evidence suggesting that V1 is involved in higher levels of visual function than is commonly assumed (e.g., Alink et al., 2010; Astorga et al., 2022; Ayzenshtat et al., 2012; Chadwick et al., 2022; Kazandjian et al., 2021; Koivisto et al., 2011; Kok & de Lange, 2014; Lamme, 1995; Lamme et al., 1998; Muckli, 2010; Noudoost & Moore, 2011; Rademaker et al., 2019; Roland et al., 2006; Super et al., 2001; Zipser et al., 1996). Because the retinotopic format of V1 coding is shared across processing levels, it offers a common platform for bottom-up and top-down communication (see Deco & Lee, 2004; Kravitz et al., 2013; Mumford, 1992). Yet, the finding that V1 is recruited for the coarse-to-fine processing of the human face is compelling. Indeed, human face perception is well documented for its reliance on high-level visual specialized neural populations (e.g., Bernstein et al., 2018; Kanwisher et al., 1997; Rossion et al., 2003b; Tsao et al., 2006). Consequently, the potential contribution of V1 to human face processing has been largely overlooked (e.g., recent Collins & Olson, 2014; Fox et al., 2009; Gobbini & Haxby, 2007; Ishai, 2008; with some noteable exceptions, e.g.: Grill-Spector et al., 2017; Petro et al., 2013). *A fortiori*, the coarse-to-fine dynamics found in V1 during face encoding indicate that the primary visual cortex acts as a generic blackboard for the recurrent refinement of representations in the human visual system (Bullier, 2001; Kok & de Lange, 2014; Lee et al., 1998). We thus believe that the present findings yield important insights for object recognition in general.

By combining backward masking with fMRI, we present evidence that the integration of visual information operates in a coarse-to-fine manner in V1. Disrupting the processing of coarse input structure disrupted V1 activity most severely in the earliest time window, and decreased in influence over time; the opposite was observed when masking HSF. Similar signatures of coarse-to-fine integration were found in fusiform, parietal and frontal regions. The robust coarse-to-fine dynamics observed in V1 indicate its active involvement in the progressive sharpening of visual representations in the human brain. The fact that V1 response dynamics to strictly identical stimulus sets differed depending on the masked scale adds to growing evidence that the role of V1 goes beyond the early processing and passive transmission of visual information to the rest of the brain. It instead indicates that V1 may yield a ‘spatially registered common forum’ or ‘blackboard’ where top-down inferences are integrated with incoming visual signals through recurrent interaction with high-level regions located in the inferotemporal, dorsal and frontal regions (Bullier, 2001; Deco & Lee, 2004; Roland, 2010).

## Acknowledgements

This work was supported by the FNRS ASP grants - 34764605 and 40005000 - awarded to JS, the Excellence of Science grant HUMVISCAT- 30991544 - and the Projet de Recherche grant - 33651302 - awarded to VG. VG is an F.R.S.-F.N.R.S. Research Associate.

## Notes

https://osf.io/echgr/

### Competing Interest Statement

The authors have declared no competing interest.

## References

Ahmed, B., Hanazawa, A., Undeman, C., Eriksson, D., Valentiniene, S., & Roland, P. E. (2008). Cortical dynamics subserving visual apparent motion. Cereb Cortex, 18(12), 2796–2810. doi:10.1093/cercor/bhn038

Ales, J. M., Farzin, F., Rossion, B., & Norcia, A. M. (2012). An objective method for measuring face detection thresholds using the sweep steady-state visual evoked response. J Vis, 12(10). doi:10.1167/12.10.18

Alink, A., Schwiedrzik, C. M., Kohler, A., Singer, W., & Muckli, L. (2010). Stimulus predictability reduces responses in primary visual cortex. J Neurosci, 30(8), 2960–2966. doi:10.1523/JNEUROSCI.3730-10.2010

Allen, E. A., & Freeman, R. D. (2006). Dynamic spatial processing originates in early visual pathways. J Neurosci, 26(45), 11763–11774. doi:10.1523/JNEUROSCI.3297-06.2006

Astorga, G., Chen, M., Yan, Y., Altavini, T. S., Jiang, C. S., Li, W., & Gilbert, C. (2022). Adaptive processing and perceptual learning in visual cortical areas V1 and V4. Proc Natl Acad Sci U S A, 119(42), e2213080119. doi:10.1073/pnas.2213080119

Ayzenberg, V., & Behrmann, M. (2022). Does the brain’s ventral visual pathway compute object shape? Trends in Cognitive Sciences.

Ayzenberg, V., Simmons, C., & Behrmann, M. (2022). Temporal asymmetries and interactions between dorsal and ventral visual pathways during object recognition. BioRxiv.

Ayzenshtat, I., Gilad, A., Zurawel, G., & Slovin, H. (2012). Population response to natural images in the primary visual cortex encodes local stimulus attributes and perceptual processing. J Neurosci, 32(40), 13971–13986. doi:10.1523/JNEUROSCI.1596-12.2012

Bacon-Mace, N., Mace, M. J., Fabre-Thorpe, M., & Thorpe, S. J. (2005). The time course of visual processing: backward masking and natural scene categorisation. Vision Res, 45(11), 1459–1469. doi:10.1016/j.visres.2005.01.004

Bar, M. (2004). Visual objects in context. Nat Rev Neurosci, 5(8), 617–629. doi:10.1038/nrn1476

Bar, M., Kassam, K. S., Ghuman, A. S., Boshyan, J., Schmid, A. M., Dale, A. M., … Halgren, E. (2006). Top-down facilitation of visual recognition. Proc Natl Acad Sci U S A, 103(2), 449–454. doi:10.1073/pnas.0507062103

Barlow, H. B. (1975). Visual experience and cortical development. Nature, 258(5532), 199–204. doi:10.1038/258199a0

Baseler, H. A., & Sutter, E. E. (1997). M and P components of the VEP and their visual field distribution. Vision Res, 37(6), 675–690. doi:10.1016/s0042-6989(96)00209-x

Bernstein, M., Erez, Y., Blank, I., & Yovel, G. (2018). An Integrated Neural Framework for Dynamic and Static Face Processing. Sci Rep, 8(1), 7036. doi:10.1038/s41598-018-25405-9

Bex, P. J., & Makous, W. (2002). Spatial frequency, phase, and the contrast of natural images. J Opt Soc Am A Opt Image Sci Vis, 19(6), 1096–1106. doi:10.1364/josaa.19.001096

Bredfeldt, C. E., & Ringach, D. L. (2002). Dynamics of spatial frequency tuning in macaque V1. J Neurosci, 22(5), 1976–1984. doi:10.1523/JNEUROSCI.22-05-01976.2002

Breitmeyer, B. G. (1975). Simple reaction time as a measure of the temporal response properties of transient and sustained channels. Vision Res, 15(12), 1411–1412. doi:10.1016/0042-6989(75)90200-x

Breitmeyer, B. G. (1984). Visual Masking: An Integrative Approach.. New York: Oxford University Press.

Budd, J. M. (1998). Extrastriate feedback to primary visual cortex in primates: a quantitative analysis of connectivity. Proc Biol Sci, 265(1400), 1037–1044. doi:10.1098/rspb.1998.0396

Bullier, J. (2001). Integrated model of visual processing. Brain Res Brain Res Rev, 36(2-3), 96–107. doi:10.1016/s0165-0173(01)00085-6

Chadwick, A., Khan, A. G., Poort, J., Blot, A., Hofer, S. B., Mrsic-Flogel, T. D., & Sahani, M. (2022). Learning shapes cortical dynamics to enhance integration of relevant sensory input. Neuron. doi:10.1016/j.neuron.2022.10.001

Collins, J. A., & Olson, I. R. (2014). Beyond the FFA: The role of the ventral anterior temporal lobes in face processing. Neuropsychologia, 61, 65–79. doi:10.1016/j.neuropsychologia.2014.06.005

Cox, R. W., & Hyde, J. S. (1997). Software tools for analysis and visualization of fMRI data. NMR Biomed, 10(4-5), 171–178. doi:10.1002/(sici)1099-1492(199706/08)10:4/5<171::aid-nbm453>3.0.co;2-l

Dale, A. M., Fischl, B., & Sereno, M. I. (1999). Cortical surface-based analysis. I. Segmentation and surface reconstruction. Neuroimage, 9(2), 179–194. doi:10.1006/nimg.1998.0395

de-Wit, L. H., Kubilius, J., Wagemans, J., & Op de Beeck, H. P. (2012). Bistable Gestalts reduce activity in the whole of V1, not just the retinotopically predicted parts. J Vis, 12(11). doi:10.1167/12.11.12

Deco, G., & Lee, T. S. (2004). The role of early visual cortex in visual integration: a neural model of recurrent interaction. Eur J Neurosci, 20(4), 1089–1100. doi:10.1111/j.1460-9568.2004.03528.x

Douglas, R. J., & Martin, K. A. (2007). Mapping the matrix: the ways of neocortex. Neuron, 56(2), 226–238. doi:10.1016/j.neuron.2007.10.017

Einevoll, G. T., Jurkus, P., & Heggelund, P. (2011). Coarse-to-fine changes of receptive fields in lateral geniculate nucleus have a transient and a sustained component that depend on distinct mechanisms. PLoS One, 6(9), e24523. doi:10.1371/journal.pone.0024523

Eriksson, D., & Roland, P. (2006). Feed-forward, feedback and lateral interactions in membrane potentials and spike trains from the visual cortex in vivo. J Physiol Paris, 100(1-3), 100–109. doi:10.1016/j.jphysparis.2006.09.009

Esteban, O., Blair, R., Markiewicz, C. J., Berleant, S. L., Moodie, C., Ma, F., & Isik, I. A. (2018). FMRIprep: Zenodo.

Esteban, O., Markiewicz, C. J., Blair, R. W., Moodie, C. A., Isik, A. I., Erramuzpe, A., … Gorgolewski, K. J. (2019). fMRIPrep: a robust preprocessing pipeline for functional MRI. Nat Methods, 16(1), 111–116. doi:10.1038/s41592-018-0235-4

Fahrenfort, J. J., Scholte, H. S., & Lamme, V. A. (2007). Masking disrupts reentrant processing in human visual cortex. J Cogn Neurosci, 19(9), 1488–1497. doi:10.1162/jocn.2007.19.9.1488

Fahrenfort, J. J., Snijders, T. M., Heinen, K., van Gaal, S., Scholte, H. S., & Lamme, V. A. (2012). Neuronal integration in visual cortex elevates face category tuning to conscious face perception. Proc Natl Acad Sci U S A, 109(52), 21504–21509. doi:10.1073/pnas.1207414110

Fintzi, A. R., & Mahon, B. Z. (2014). A bimodal tuning curve for spatial frequency across left and right human orbital frontal cortex during object recognition. Cereb Cortex, 24(5), 1311–1318. doi:10.1093/cercor/bhs419

Finzi, D., Gomez, J., Nordt, M., Rezai, A. A., Poltoratski, S., & Grill-Spector, K. (2021). Differential spatial computations in ventral and lateral face-selective regions are scaffolded by structural connections. Nat Commun, 12(1), 2278. doi:10.1038/s41467-021-22524-2

Fox, C. J., Iaria, G., & Barton, J. J. (2009). Defining the face processing network: optimization of the functional localizer in fMRI. Hum Brain Mapp, 30(5), 1637–1651. doi:10.1002/hbm.20630

Frazor, R. A., Albrecht, D. G., Geisler, W. S., & Crane, A. M. (2004). Visual cortex neurons of monkeys and cats: temporal dynamics of the spatial frequency response function. J Neurophysiol, 91(6), 2607–2627. doi:10.1152/jn.00858.2003

Freud, E., Culham, J. C., Plaut, D. C., & Behrmann, M. (2017). The large-scale organization of shape processing in the ventral and dorsal pathways. Elife, 6. doi:10.7554/eLife.27576

Gao, Z., & Bentin, S. (2011). Coarse-to-fine encoding of spatial frequency information into visual short-term memory for faces but impartial decay. J Exp Psychol Hum Percept Perform, 37(4), 1051–1064. doi:10.1037/a0023091

Gentile, F., Ales, J., & Rossion, B. (2017). Being BOLD: The neural dynamics of face perception. Hum Brain Mapp, 38(1), 120–139. doi:10.1002/hbm.23348

George, N., Dolan, R. J., Fink, G. R., Baylis, G. C., Russell, C., & Driver, J. (1999). Contrast polarity and face recognition in the human fusiform gyrus. Nat Neurosci, 2(6), 574–580. doi:10.1038/9230

Gilbert, C. D., & Li, W. (2013). Top-down influences on visual processing. Nat Rev Neurosci, 14(5), 350–363. doi:10.1038/nrn3476

Gobbini, M. I., & Haxby, J. V. (2007). Neural systems for recognition of familiar faces. Neuropsychologia, 45(1), 32–41. doi:10.1016/j.neuropsychologia.2006.04.015

Goffaux, V., Duecker, F., Hausfeld, L., Schiltz, C., & Goebel, R. (2016). Horizontal tuning for faces originates in high-level Fusiform Face Area. Neuropsychologia, 81, 1–11. doi:10.1016/j.neuropsychologia.2015.12.004

Goffaux, V., Peters, J., Haubrechts, J., Schiltz, C., Jansma, B., & Goebel, R. (2011). From coarse to fine? Spatial and temporal dynamics of cortical face processing. Cereb Cortex, 21(2), 467–476. doi:10.1093/cercor/bhq112

Goffaux, V., Schiltz, C., Mur, M., & Goebel, R. (2012). Local discriminability determines the strength of holistic processing for faces in the fusiform face area. Front Psychol, 3, 604. doi:10.3389/fpsyg.2012.00604

Gorgolewski, K., Burns, C. D., Madison, C., Clark, D., Halchenko, Y. O., Waskom, M. L., & Ghosh, S. S. (2011). Nipype: a flexible, lightweight and extensible neuroimaging data processing framework in python. Front Neuroinform, 5, 13. doi:10.3389/fninf.2011.00013

Gorgolewski, K. J., Auer, T., Calhoun, V. D., Craddock, R. C., Das, S., Duff, E. P., … Poldrack, R. A. (2016). The brain imaging data structure, a format for organizing and describing outputs of neuroimaging experiments. Sci Data, 3, 160044. doi:10.1038/sdata.2016.44

Gorgolewski, K. J., Nichols, T., Kennedy, D. N., Poline, J. B., & Poldrack, R. A. (2018). Making replication prestigious. Behav Brain Sci, 41, e131. doi:10.1017/S0140525X18000663

Green, M. F., Glahn, D., Engel, S. A., Nuechterlein, K. H., Sabb, F., Strojwas, M., & Cohen, M. S. (2005). Regional brain activity associated with visual backward masking. J Cogn Neurosci, 17(1), 13–23. doi:10.1162/0898929052880011

Greve, D. N., & Fischl, B. (2009). Accurate and robust brain image alignment using boundary-based registration. Neuroimage, 48(1), 63–72. doi:10.1016/j.neuroimage.2009.06.060

Grill-Spector, K., Kushnir, T., Hendler, T., & Malach, R. (2000). The dynamics of object-selective activation correlate with recognition performance in humans. Nat Neurosci, 3(8), 837–843. doi:10.1038/77754

Grill-Spector, K., Weiner, K. S., Kay, K., & Gomez, J. (2017). The Functional Neuroanatomy of Human Face Perception. Annu Rev Vis Sci, 3, 167–196. doi:10.1146/annurev-vision-102016-061214

Halit, H., de Haan, M., Schyns, P. G., & Johnson, M. H. (2006). Is high-spatial frequency information used in the early stages of face detection? Brain Res, 1117(1), 154–161. doi:10.1016/j.brainres.2006.07.059

Hansen, B. C., Greene, M. R., & Field, D. J. (2021). Dynamic Electrode-to-Image (DETI) mapping reveals the human brain’s spatiotemporal code of visual information. PLoS Comput Biol, 17(9), e1009456. doi:10.1371/journal.pcbi.1009456

Harvey, M. A., Valentiniene, S., & Roland, P. E. (2009). Cortical Membrane Potential Dynamics and Laminar Firing during Object Motion. Front Syst Neurosci, 3, 7. doi:10.3389/neuro.06.007.2009

Haynes, J. D., Driver, J., & Rees, G. (2005). Visibility reflects dynamic changes of effective connectivity between V1 and fusiform cortex. Neuron, 46(5), 811–821. doi:10.1016/j.neuron.2005.05.012

Hegde, J. (2008). Time course of visual perception: coarse-to-fine processing and beyond. Prog Neurobiol, 84(4), 405–439. doi:10.1016/j.pneurobio.2007.09.001

Heinze, G., Wallisch, C., & Dunkler, D. (2018). Variable selection - A review and recommendations for the practicing statistician. Biom J, 60(3), 431–449. doi:10.1002/bimj.201700067

Hughes, H. C., Nozawa, G., & Kitterle, F. (1996). Global precedence, spatial frequency channels, and the statistics of natural images. J Cogn Neurosci, 8(3), 197–230. doi:10.1162/jocn.1996.8.3.197

Hupe, J. M., James, A. C., Payne, B. R., Lomber, S. G., Girard, P., & Bullier, J. (1998). Cortical feedback improves discrimination between figure and background by V1, V2 and V3 neurons. Nature, 394(6695), 784–787. doi:10.1038/29537

Ishai, A. (2008). Let’s face it: it’s a cortical network. Neuroimage, 40(2), 415–419. doi:10.1016/j.neuroimage.2007.10.040

Jacobs, C., Petras, K., Moors, P., & Goffaux, V. (2020). Contrast versus identity encoding in the face image follow distinct orientation selectivity profiles. PLoS One, 15(3), e0229185. doi:10.1371/journal.pone.0229185

Jenkinson, M., Bannister, P., Brady, M., & Smith, S. (2002). Improved optimization for the robust and accurate linear registration and motion correction of brain images. Neuroimage, 17(2), 825–841. doi:10.1016/s1053-8119(02)91132-8

Jones, R., & Keck, M. J. (1978). Visual evoked response as a function of grating spatial frequency. Invest Ophthalmol Vis Sci, 17(7), 652–659.

Kanwisher, N., McDermott, J., & Chun, M. M. (1997). The fusiform face area: a module in human extrastriate cortex specialized for face perception. J Neurosci, 17(11), 4302–4311.

Kar, K., & DiCarlo, J. J. (2021). Fast Recurrent Processing via Ventrolateral Prefrontal Cortex Is Needed by the Primate Ventral Stream for Robust Core Visual Object Recognition. Neuron, 109(1), 164–176 e165. doi:10.1016/j.neuron.2020.09.035

Kar, K., Kubilius, J., Schmidt, K., Issa, E. B., & DiCarlo, J. J. (2019). Evidence that recurrent circuits are critical to the ventral stream’s execution of core object recognition behavior. Nat Neurosci, 22(6), 974–983. doi:10.1038/s41593-019-0392-5

Kauffmann, L., Bourgin, J., Guyader, N., & Peyrin, C. (2015a). The Neural Bases of the Semantic Interference of Spatial Frequency-based Information in Scenes. J Cogn Neurosci, 27(12), 2394–2405. doi:10.1162/jocn_a_00861

Kauffmann, L., Chauvin, A., Guyader, N., & Peyrin, C. (2015b). Rapid scene categorization: role of spatial frequency order, accumulation mode and luminance contrast. Vision Res, 107, 49–57. doi:10.1016/j.visres.2014.11.013

Kauffmann, L., Chauvin, A., Pichat, C., & Peyrin, C. (2015c). Effective connectivity in the neural network underlying coarse-to-fine categorization of visual scenes. A dynamic causal modeling study. Brain Cogn, 99, 46–56. doi:10.1016/j.bandc.2015.07.004

Kauffmann, L., Ramanoel, S., Guyader, N., Chauvin, A., & Peyrin, C. (2015d). Spatial frequency processing in scene-selective cortical regions. Neuroimage, 112, 86–95.doi:10.1016/j.neuroimage.2015.02.058

Kay, K. N., & Yeatman, J. D. (2016). Bottom-up and top-down computations in high-level visual cortex. BioRxiv, 053595.

Kazandjian, T. D., Petras, D., Robinson, S. D., van Thiel, J., Greene, H. W., Arbuckle, K., Casewell, N.R. (2021). Convergent evolution of pain-inducing defensive venom components in spitting cobras. Science, 371(6527), 386–390. doi:10.1126/science.abb9303

Klein, A., Ghosh, S. S., Bao, F. S., Giard, J., Hame, Y., Stavsky, E., Keshavan, A. (2017). Mindboggling morphometry of human brains. PLoS Comput Biol, 13(2), e1005350. doi:10.1371/journal.pcbi.1005350

Koivisto, M., Railo, H., Revonsuo, A., Vanni, S., & Salminen-Vaparanta, N. (2011). Recurrent processing in V1/V2 contributes to categorization of natural scenes. J Neurosci, 31(7), 2488–2492. doi:10.1523/JNEUROSCI.3074-10.2011

Kok, P., & de Lange, F. P. (2014). Shape perception simultaneously up- and downregulates neural activity in the primary visual cortex. Curr Biol, 24(13), 1531–1535. doi:10.1016/j.cub.2014.05.042

Kovacs, G., Vogels, R., & Orban, G. A. (1995). Cortical correlate of pattern backward masking. Proc Natl Acad Sci U S A, 92(12), 5587–5591. doi:10.1073/pnas.92.12.5587

Kravitz, D. J., Saleem, K. S., Baker, C. I., Ungerleider, L. G., & Mishkin, M. (2013). The ventral visual pathway: an expanded neural framework for the processing of object quality. Trends Cogn Sci, 17(1), 26–49. doi:10.1016/j.tics.2012.10.011

Kreiman, G., & Serre, T. (2020). Beyond the feedforward sweep: feedback computations in the visual cortex. Ann N Y Acad Sci, 1464(1), 222–241. doi:10.1111/nyas.14320

Kveraga, K., Boshyan, J., & Bar, M. (2007). Magnocellular projections as the trigger of top-down facilitation in recognition. J Neurosci, 27(48), 13232–13240. doi:10.1523/JNEUROSCI.3481-07.2007

Laguesse, R., Dormal, G., Biervoye, A., Kuefner, D., & Rossion, B. (2012). Extensive visual training in adulthood significantly reduces the face inversion effect. J Vis, 12(10), 14. doi:10.1167/12.10.14

Lamme, V. A. (1995). The neurophysiology of figure-ground segregation in primary visual cortex. J Neurosci, 15(2), 1605–1615. doi:10.1523/JNEUROSCI.15-02-01605.1995

Lamme, V. A., Super, H., & Spekreijse, H. (1998). Feedforward, horizontal, and feedback processing in the visual cortex. Curr Opin Neurobiol, 8(4), 529–535. doi:10.1016/s0959-4388(98)80042-1

Lamme, V. A., Zipser, K., & Spekreijse, H. (2002). Masking interrupts figure-ground signals in V1. J Cogn Neurosci, 14(7), 1044–1053. doi:10.1162/089892902320474490

Lee, T. S. (2015). The visual system’s internal model of the world. Proc IEEE Inst Electr Electron Eng, 103(8), 1359–1378. doi:10.1109/JPROC.2015.2434601

Lee, T. S., Mumford, D., Romero, R., & Lamme, V. A. (1998). The role of the primary visual cortex in higher level vision. Vision Res, 38(15-16), 2429–2454. doi:10.1016/s0042-6989(97)00464-1

Lerner, Y., Hendler, T., Ben-Bashat, D., Harel, M., & Malach, R. (2001). A hierarchical axis of object processing stages in the human visual cortex. Cereb Cortex, 11(4), 287–297. doi:10.1093/cercor/11.4.287

Liu-Shuang, J., Yang, Y. F., Rossion, B., & Goffaux, V. (2022). Natural contrast statistics facilitate human face categorization.

Loftus, G. R., & Masson, M. E. (1994). Using confidence intervals in within-subject designs. Psychon Bull Rev, 1(4), 476–490. doi:10.3758/BF03210951

Lu, Y., Yin, J., Chen, Z., Gong, H., Liu, Y., Qian, L., … Wang, W. (2018). Revealing Detail along the Visual Hierarchy: Neural Clustering Preserves Acuity from V1 to V4. Neuron, 98(2), 417–428 e413. doi:10.1016/j.neuron.2018.03.009

Malone, B. J., Kumar, V. R., & Ringach, D. L. (2007). Dynamics of receptive field size in primary visual cortex. J Neurophysiol, 97(1), 407–414. doi:10.1152/jn.00830.2006

Markov, N. T., Vezoli, J., Chameau, P., Falchier, A., Quilodran, R., Huissoud, C., … Kennedy, H. (2014). Anatomy of hierarchy: feedforward and feedback pathways in macaque visual cortex. J Comp Neurol, 522(1), 225–259. doi:10.1002/cne.23458

Mazer, J. A., Vinje, W. E., McDermott, J., Schiller, P. H., & Gallant, J. L. (2002). Spatial frequency and orientation tuning dynamics in area V1. Proc Natl Acad Sci U S A, 99(3), 1645–1650. doi:10.1073/pnas.022638499

Merkel, D. (2014). Docker: lightweight linux containers for consistent development and deployment. Linux Journal, 239(2).

Mihaylova, M., Stomonyakov, V., & Vassilev, A. (1999). Peripheral and central delay in processing high spatial frequencies: reaction time and VEP latency studies. Vision Res, 39(4), 699–705. doi:10.1016/s0042-6989(98)00165-5

Mohsenzadeh, Y., Qin, S., Cichy, R. M., & Pantazis, D. (2018). Ultra-Rapid serial visual presentation reveals dynamics of feedforward and feedback processes in the ventral visual pathway. Elife, 7. doi:10.7554/eLife.36329

Muckli, L. (2010). What are we missing here? Brain imaging evidence for higher cognitive functions in primary visual cortex V1.. International Journal of Imaging Systems and Technology, 20(2), 131–139.

Muckli, L., & Petro, L. S. (2013). Network interactions: non-geniculate input to V1. Curr Opin Neurobiol, 23(2), 195–201. doi:10.1016/j.conb.2013.01.020

Mumford, D. (1992). On the computational architecture of the neocortex. II. The role of cortico-cortical loops. Biol Cybern, 66(3), 241–251. doi:10.1007/BF00198477

Musel, B., Kauffmann, L., Ramanoel, S., Giavarini, C., Guyader, N., Chauvin, A., & Peyrin, C. (2014). Coarse-to-fine categorization of visual scenes in scene-selective cortex. J Cogn Neurosci, 26(10), 2287–2297. doi:10.1162/jocn_a_00643

Musselwhite, M. J., & Jeffreys, D. A. (1985). The influence of spatial frequency on the reaction times and evoked potentials recorded to grating pattern stimuli. Vision Res, 25(11), 1545–1555. doi:10.1016/0042-6989(85)90125-7

Nirody, J. A. (2014). Development of spatial coarse-to-fine processing in the visual pathway. J Comput Neurosci, 36(3), 401–414. doi:10.1007/s10827-013-0480-6

Noudoost, B., & Moore, T. (2011). Control of visual cortical signals by prefrontal dopamine. Nature, 474(7351), 372–375. doi:10.1038/nature09995

Nowak, L. G., Munk, M. H., Girard, P., & Bullier, J. (1995). Visual latencies in areas V1 and V2 of the macaque monkey. Vis Neurosci, 12(2), 371–384. doi:10.1017/s095252380000804x

O’Reilly, J. X., Woolrich, M. W., Behrens, T. E., Smith, S. M., & Johansen-Berg, H. (2012). Tools of the trade: psychophysiological interactions and functional connectivity. Soc Cogn Affect Neurosci, 7(5), 604–609. doi:10.1093/scan/nss055

Oldfield, R. C. (1971). The assessment and analysis of handedness: the Edinburgh inventory. Neuropsychologia, 9(1), 97–113. doi:10.1016/0028-3932(71)90067-4

Olshausen, B. A., & Field, D. J. (1996). Emergence of simple-cell receptive field properties by learning a sparse code for natural images. Nature, 381(6583), 607–609. doi:10.1038/381607a0

Olshausen, B. A., & Field, J. D. (2006). What is the other 85 percent of V1 doing? In J. L. Van Hemmen & T. J. Sejnowski (Eds.), 23 Problems in Systems Neuroscience (pp. 182–212). New York: Oxford University Press.

Parker, D. M. (1980). Simple reaction times to the onset, offset, and contrast reversal of sinusoidal grating stimuli. Percept Psychophys, 28(4), 365–368. doi:10.3758/bf03204396

Parker, D. M., & Dutch, S. (1987). Perceptual latency and spatial frequency. Vision Res, 27(8), 1279–1283. doi:10.1016/0042-6989(87)90204-5

Parker, D. M., Lishman, J. R., & Hughes, J. (1992). Temporal integration of spatially filtered visual images. Perception, 21(2), 147–160. doi:10.1068/p210147

Parker, D. M., Lishman, J. R., & Hughes, J. (1997). Evidence for the view that temporospatial integration in vision is temporally anisotropic. Perception, 26(9), 1169–1180. doi:10.1068/p261169

Parker, D. M., & Salzen, E. A. (1977). The spatial selectivity of early and late waves within the human visual evoked response. Perception, 6(1), 85–95. doi:10.1068/p060085

Petras, K., Ten Oever, S., Dalal, S. S., & Goffaux, V. (2021). Information redundancy across spatial scales modulates early visual cortical processing. Neuroimage, 244, 118613. doi:10.1016/j.neuroimage.2021.118613

Petras, K., Ten Oever, S., Jacobs, C., & Goffaux, V. (2019). Coarse-to-fine information integration in human vision. Neuroimage, 186, 103–112. doi:10.1016/j.neuroimage.2018.10.086

Petro, L. S., Smith, F. W., Schyns, P. G., & Muckli, L. (2013). Decoding face categories in diagnostic subregions of primary visual cortex. Eur J Neurosci, 37(7), 1130–1139. doi:10.1111/ejn.12129

Petro, L. S., Vizioli, L., & Muckli, L. (2014). Contributions of cortical feedback to sensory processing in primary visual cortex. Front Psychol, 5, 1223. doi:10.3389/fpsyg.2014.01223

Peyrin, C., Mermillod, M., Chokron, S., & Marendaz, C. (2006). Effect of temporal constraints on hemispheric asymmetries during spatial frequency processing. Brain Cogn, 62(3), 214–220. doi:10.1016/j.bandc.2006.05.005

Peyrin, C., Michel, C. M., Schwartz, S., Thut, G., Seghier, M., Landis, T., … Vuilleumier, P. (2010). The neural substrates and timing of top-down processes during coarse-to-fine categorization of visual scenes: a combined fMRI and ERP study. J Cogn Neurosci, 22(12), 2768–2780. doi:10.1162/jocn.2010.21424

Power, J. D., Mitra, A., Laumann, T. O., Snyder, A. Z., Schlaggar, B. L., & Petersen, S. E. (2014). Methods to detect, characterize, and remove motion artifact in resting state fMRI. Neuroimage, 84, 320–341. doi:10.1016/j.neuroimage.2013.08.048

Rademaker, R. L., Chunharas, C., & Serences, J. T. (2019). Coexisting representations of sensory and mnemonic information in human visual cortex. Nat Neurosci, 22(8), 1336–1344. doi:10.1038/s41593-019-0428-x

Rieger, J. W., Gegenfurtner, K. R., Schalk, F., Koechy, N., Heinze, H. J., & Grueschow, M. (2013). BOLD responses in human V1 to local structure in natural scenes: Implications for theories of visual coding. J Vis, 13(2), 19. doi:10.1167/13.2.19

Roland, P. E. (2010). Six principles of visual cortical dynamics. Front Syst Neurosci, 4, 28. doi:10.3389/fnsys.2010.00028

Roland, P. E., Hanazawa, A., Undeman, C., Eriksson, D., Tompa, T., Nakamura, H., … Ahmed, B. (2006). Cortical feedback depolarization waves: a mechanism of top-down influence on early visual areas. Proc Natl Acad Sci U S A, 103(33), 12586–12591. doi:10.1073/pnas.0604925103

Rolls, E. T., & Tovee, M. J. (1994). Processing speed in the cerebral cortex and the neurophysiology of visual masking. Proc Biol Sci, 257(1348), 9–15. doi:10.1098/rspb.1994.0087

Rolls, E. T., Tovee, M. J., & Panzeri, S. (1999). The neurophysiology of backward visual masking: information analysis. J Cogn Neurosci, 11(3), 300–311. doi:10.1162/089892999563409

Rossion, B., Caldara, R., Seghier, M., Schuller, A. M., Lazeyras, F., & Mayer, E. (2003a). A network of occipito-temporal face-sensitive areas besides the right middle fusiform gyrus is necessary for normal face processing. Brain, 126(Pt 11), 2381–2395. doi:10.1093/brain/awg241

Rossion, B., Dricot, L., Devolder, A., Bodart, J. M., Crommelinck, M., De Gelder, B., & Zoontjes, R. (2000). Hemispheric asymmetries for whole-based and part-based face processing in the human fusiform gyrus. J Cogn Neurosci, 12(5), 793–802. doi:10.1162/089892900562606

Rossion, B., Hanseeuw, B., & Dricot, L. (2012). Defining face perception areas in the human brain: a large-scale factorial fMRI face localizer analysis. Brain Cogn, 79(2), 138–157. doi:10.1016/j.bandc.2012.01.001

Rossion, B., Schiltz, C., & Crommelinck, M. (2003b). The functionally defined right occipital and fusiform “face areas” discriminate novel from visually familiar faces. Neuroimage, 19(3), 877–883. doi:10.1016/s1053-8119(03)00105-8

Schiltz, C., & Rossion, B. (2006). Faces are represented holistically in the human occipito-temporal cortex. Neuroimage, 32(3), 1385–1394. doi:10.1016/j.neuroimage.2006.05.037

Schmolesky, M. T., Wang, Y., Hanes, D. P., Thompson, K. G., Leutgeb, S., Schall, J. D., & Leventhal, A. G. (1998). Signal timing across the macaque visual system. J Neurophysiol, 79(6), 3272–3278. doi:10.1152/jn.1998.79.6.3272

Scholte, H. S., Jolij, J., Fahrenfort, J. J., & Lamme, V. A. (2008). Feedforward and recurrent processing in scene segmentation: electroencephalography and functional magnetic resonance imaging. J Cogn Neurosci, 20(11), 2097–2109. doi:10.1162/jocn.2008.20142

Schyns, & Oliva, A. (1994). Evidence for time- and spatial-scale-dependent scene recognition. Psychological Science, 5, 195–201.

Stigliani, A., Weiner, K. S., & Grill-Spector, K. (2015). Temporal Processing Capacity in High-Level Visual Cortex Is Domain Specific. J Neurosci, 35(36), 12412–12424. doi:10.1523/JNEUROSCI.4822-14.2015

Stromeyer, C. F., 3rd, & Julesz, B. (1972). Spatial-frequency masking in vision: critical bands and spread of masking. J Opt Soc Am, 62(10), 1221–1232. doi:10.1364/josa.62.001221

Super, H., Spekreijse, H., & Lamme, V. A. (2001). A neural correlate of working memory in the monkey primary visual cortex. Science, 293(5527), 120–124. doi:10.1126/science.1060496

Takagaki, K., Zhang, C., Wu, J. Y., & Lippert, M. T. (2008). Crossmodal propagation of sensory-evoked and spontaneous activity in the rat neocortex. Neurosci Lett, 431(3), 191–196. doi:10.1016/j.neulet.2007.11.069

Tanaka, H., & Sawada, R. (2022). Dynamics and Mechanisms of Contrast-Dependent Modulation of Spatial-Frequency Tuning in the Early Visual Cortex. J Neurosci, 42(37), 7047–7059. doi:10.1523/JNEUROSCI.2086-21.2022

Thome, I., Hohmann, D. M., Zimmermann, K. M., Smith, M. L., Kessler, R., & Jansen, A. (2021). “I Spy with my Little Eye, Something that is a Face…”: A Brain Network for Illusory Face Detection. Cereb Cortex, 32(1), 137–157. doi:10.1093/cercor/bhab199

Tsao, D. Y., Freiwald, W. A., Tootell, R. B., & Livingstone, M. S. (2006). A cortical region consisting entirely of face-selective cells. Science, 311(5761), 670–674. doi:10.1126/science.1119983

Tustison, N. J., Avants, B. B., Cook, P. A., Zheng, Y., Egan, A., Yushkevich, P. A., & Gee, J. C. (2010). N4ITK: improved N3 bias correction. IEEE Trans Med Imaging, 29(6), 1310–1320. doi:10.1109/TMI.2010.2046908

Van Dromme, I. C., Premereur, E., Verhoef, B. E., Vanduffel, W., & Janssen, P. (2016). Posterior Parietal Cortex Drives Inferotemporal Activations During Three-Dimensional Object Vision. PLoS Biol, 14(4), e1002445. doi:10.1371/journal.pbio.1002445

Vlamings, P. H., Goffaux, V., & Kemner, C. (2009). Is the early modulation of brain activity by fearful facial expressions primarily mediated by coarse low spatial frequency information? J Vis, 9(5), 12 11–13. doi:10.1167/9.5.12

Watt, R. J. (1987). Scanning from coarse to fine spatial scales in the human visual system after the onset of a stimulus. J Opt Soc Am A, 4(10), 2006–2021. doi:10.1364/josaa.4.002006

Weiner, K. S., & Grill-Spector, K. (2013). Neural representations of faces and limbs neighbor in human high-level visual cortex: evidence for a new organization principle. Psychol Res, 77(1), 74–97. doi:10.1007/s00426-011-0392-x

Wibral, M., Bledowski, C., Kohler, A., Singer, W., & Muckli, L. (2009). The timing of feedback to early visual cortex in the perception of long-range apparent motion. Cereb Cortex, 19(7), 1567–1582. doi:10.1093/cercor/bhn192

Wyatte, D., Jilk, D. J., & O’Reilly, R. C. (2014). Early recurrent feedback facilitates visual object recognition under challenging conditions. Front Psychol, 5, 674. doi:10.3389/fpsyg.2014.00674

Yue, X., Nasr, S., Devaney, K. J., Holt, D. J., & Tootell, R. B. (2013). fMRI analysis of contrast polarity in face-selective cortex in humans and monkeys. Neuroimage, 76, 57–69. doi:10.1016/j.neuroimage.2013.02.068

Zachariou, V., Nikas, C. V., Safiullah, Z. N., Gotts, S. J., & Ungerleider, L. G. (2017). Spatial mechanisms within the dorsal visual pathway contribute to the configural processing of faces. Cerebral Cortex, 27(8), 4124–4138.

Zhang, Y., Brady, M., & Smith, S. (2001). Segmentation of brain MR images through a hidden Markov random field model and the expectation-maximization algorithm. IEEE Trans Med Imaging, 20(1), 45–57. doi:10.1109/42.906424

Zipser, K., Lamme, V. A., & Schiller, P. H. (1996). Contextual modulation in primary visual cortex. J Neurosci, 16(22), 7376–7389. doi:10.1523/JNEUROSCI.16-22-07376.1996

