## Supplementary material for "Backward masking reveals coarse-to-fine dynamics in human V1"

### Supplementary material 1

#### Face localiser stimuli

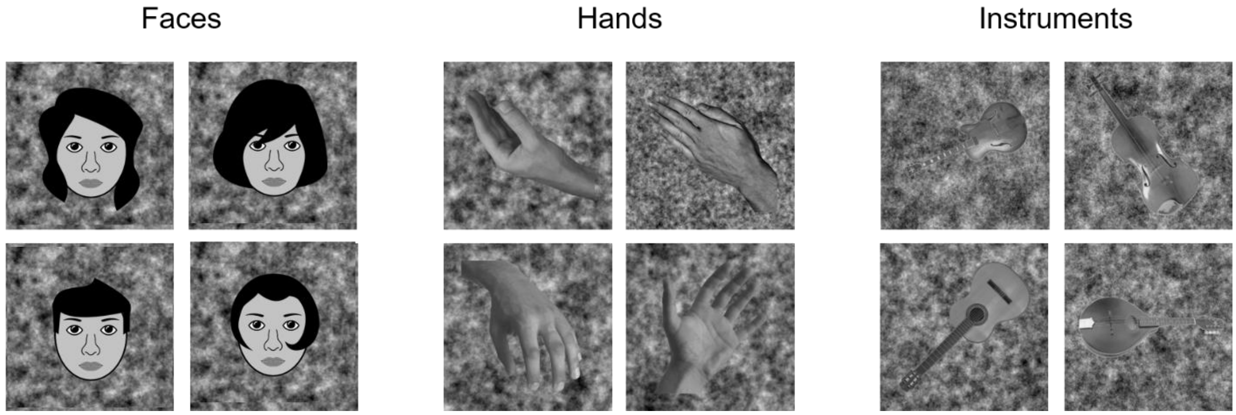


*Figure 1.* Stimuli as used in the functional localiser experiment. Stimuli were adapted from the *fLoc functional localizer package* (Stigliani et al., 2015). The background for the stimuli were created using the iterative phase scrambling method as described in the method section to ensure a uniform distribution of SF and orientation content over the whole image. Cartoon faces are used in replacement of photographs in the figure as journal regulations prohibit the use of identifiable faces.

### Supplementary material 2

#### Plots of beta values from GLM

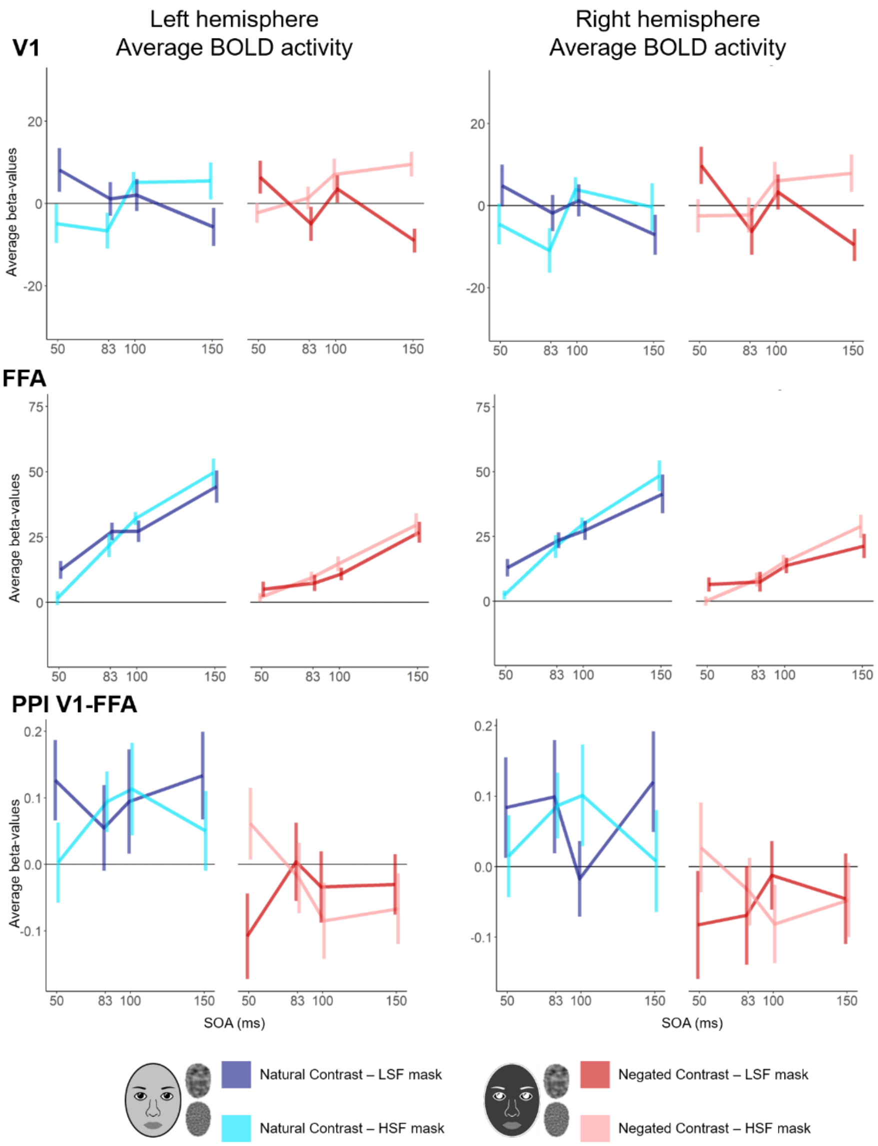


*Figure 2.* Intact-scrambled response differences for V1, FFA and for the connectivity analysis between these regions, for both left (left column) and right (right column) hemispheres. The group averaged beta-values include a 95% within-subjects confidence interval as described in Loftus & Masson, 1994. Data is just for illustration, since no hemispherical differences were found in the analyses. Cartoon faces are used in replacement of photographs in the figure as journal regulations prohibit the use of identifiable faces.

### Supplementary material 3

Table 1

*Step-by-step overview of the stepwise variable selection procedure following backward elimination (Heinze et al., 2018). The data is either accuracy of the behavioural task, ROI data (V1, V2, FFA or OFA) or the PPI between V1 and FFA. The first model is the most complex model containing all interaction terms, next the non-significant (interaction) terms were stepwise eliminated from the model.*

| *Model number* | *Model* | *BIC* |
|---|---|---|
| | Accuracy | |
| 1 | Acurracy ~ 1 + SOA * SFmask * StimulusType + (1 \| subject) | -1298.851 |
| 2 | Acurracy ~ 1 + SOA * SFmask + SOA * StimulusType + SFmask * StimulusType + (1 \| subject) | -1343.635 |
| 3 | Acurracy ~ 1 + SOA + SFmask + StimulusType + (1 \| subject) | -1443.545 |
| | V1 | |
| 1 | BetaValue ~ 1 + SOA * SFmask * StimulusType * Hemisphere + (1 \| subject) | 118743.7 |
| 2 | BetaValue ~ 1 + SOA * SFmask * StimulusType + SOA * SFmask * Hemisphere + SOA * StimulusType * Hemisphere + SFmask * StimulusType * Hemisphere + (1 \| subject) | 118729 |
| 3 | BetaValue ~ 1 + SOA * SFmask + SOA * StimulusType + SFmask * StimulusType + SOA * Hemisphere + SFmask * Hemisphere + StimulusType * Hemisphere + (1 \| subject) | 118676.6 |
| 4 | BetaValue ~ 1 + SOA * SFmask + SFmask * StimulusType + (1 \| subject) | 118624.6 |

V2

| | | |
|---|---|---|
| 1 | BetaValue ~ 1 + SOA * SFmask * StimulusType * Hemisphere + (1 \| subject) | 114363.5 |
| 2 | BetaValue ~ 1 + SOA * SFmask * StimulusType + SOA * SFmask * Hemisphere + SOA * StimulusType * Hemisphere + SFmask * StimulusType * Hemisphere + (1 \| subject) | 114348.1 |
| 3 | BetaValue ~ 1 + SOA * SFmask + SOA * StimulusType + SFmask * StimulusType + SOA * Hemisphere + SFmask * Hemisphere + StimulusType * Hemisphere + (1 \| subject) | 114294.2 |
| 4 | BetaValue ~ 1 + SOA * SFmask + SFmask * StimulusType + (1 \| subject) | 114240.2 |

FFA

| | | |
|---|---|---|
| 1 | BetaValue ~ 1 + SOA * SFmask * StimulusType * Hemisphere + (1 \| subject) | 107593.5 |
| 2 | BetaValue ~ 1 + SOA * SFmask * StimulusType + SOA * SFmask * Hemisphere + SOA * StimulusType * Hemisphere + SFmask * StimulusType * Hemisphere + (1 \| subject) | 107577.8 |
| 3 | BetaValue ~ 1 + SOA * SFmask + SOA * StimulusType + SFmask * StimulusType + SOA * Hemisphere + SFmask * Hemisphere + StimulusType * Hemisphere + (1 \| subject) | 107523.6 |
| 4 | BetaValue ~ 1 + SOA * SFmask + SOA * StimulusType + (1 \| subject) | 107475 |

| | | |
|---|---|---|
| | OFA | |
| 1 | BetaValue ~ 1 + SOA * SFmask * StimulusType * Hemisphere + (1 \| subject) | 113935.8 |
| 2 | BetaValue ~ 1 + SOA * SFmask * StimulusType + SOA * SFmask * Hemisphere + SOA * StimulusType * Hemisphere + SFmask * StimulusType * Hemisphere + (1 \| subject) | 113920.3 |
| 3 | BetaValue ~ 1 + SOA * SFmask + SOA * StimulusType + SFmask * StimulusType + SOA * Hemisphere + SFmask * Hemisphere + StimulusType * Hemisphere + (1 \| subject) | 113866.5 |
| 4 | BetaValue ~ 1 + SOA * SFmask + SOA * StimulusType + (1 \| subject) | 113824.1 |
| | PPI (V1-FFA) | |
| 1 | BetaValue ~ 1 + SOA * SFmask * StimulusType * Hemisphere + (1 \| subject) | 28709.61 |
| 2 | BetaValue ~ 1 + SOA * SFmask * StimulusType + SOA * SFmask * Hemisphere + SOA * StimulusType * Hemisphere + SFmask * StimulusType * Hemisphere + (1 \| subject) | 28692.92 |
| 3 | BetaValue ~ 1 + SFmask * StimulusType + SOA * SFmask + SOA * StimulusType + SOA * Hemisphere + SFmask * Hemisphere + StimulusType * Hemisphere + (1 \| subject) | 28623.33 |
| 4 | BetaValue ~ 1 + SOA + SFmask + Hemisphere + StimulusType + (1 \| subject) | 28529.54 |
| 5 | BetaValue ~ 1 + StimulusType + (1 \| subject) | 28482.8 |

### Supplementary material 4

Supplementary Table 2.

*Accuracy Fixed effects*

| | *Estimate* | *SE* | *df* | *T* | *p* | |
|---|---|---|---|---|---|---|
| (Intercept) | 0.97260 | 0.00487 | 225.6 | 199.824 | < 0.001 | *** |
| SOA | -0.00002 | 0.00005 | 364 | -0.542 | 0.588 | |
| SFmask | -0.00181 | 0.00164 | 364 | -1.103 | 0.271 | |
| StimulusType[natural] | 0.00215 | 0.00232 | 364 | 0.928 | 0.354 | |
| StimulusType[negated] | 0.00181 | 0.00232 | 364 | 0.778 | 0.437 | |

Supplementary Table 3.

*V1 Fixed effects*

| | *Estimate* | *SE* | *df* | *T* | *p* | |
|---|---|---|---|---|---|---|
| (Intercept) | 1.724 | 2.392 | 300.3 | 0.721 | 0.472 | |
| SOA | -0.01528 | 0.02204 | 10190 | -0.693 | 0.488 | |
| SFmask | 11.03 | 2.256 | 10190 | 4.892 | < 0.001 | *** |
| StimulusType | 0.8803 | 0.796 | 10190 | 1.106 | 0.269 | |
| SOA:SFmask | -0.1204 | 0.02204 | 10190 | -5.462 | < 0.001 | *** |
| SFmask:StimulusType | -1.466 | 0.796 | 10190 | -1.842 | 0.066 | . |

Supplementary Table 4.

*FFA Fixed effects*

| | *Estimate* | *SE* | *df* | *T* | *p* | |
|---|---|---|---|---|---|---|
| (Intercept) | -9.929 | 2.368 | 27.77 | -4.193 | < 0.001 | *** |
| SOA | 0.3092 | 0.01274 | 10190 | 24.279 | < 0.001 | *** |
| SFmask | 6.397 | 1.303 | 10190 | 4.908 | < 0.001 | *** |
| StimulusType | 0.03893 | 1.303 | 10190 | 0.03 | 0.976 | |
| SOA:SFmask | -0.06636 | 0.01274 | 10190 | -5.21 | < 0.001 | *** |
| SOA:StimulusType | -0.07075 | 0.01274 | 10190 | -5.555 | < 0.001 | *** |

Supplementary Table 5.

*PPI Fixed effects*

| | *Estimate* | *SE* | *df* | *T* | *p* | |
|---|---|---|---|---|---|---|
| (Intercept) | 0.01682 | 0.01831 | 15.02 | 0.919 | 0.373 | |
| StimulusType | -0.05591 | 0.009632 | 10190 | -5.804 | < 0.001 | *** |

Supplementary Table 6.

*V2 Fixed effects*

| | *Estimate* | *SE* | *df* | *T* | *p* | |
|---|---|---|---|---|---|---|
| (Intercept) | 0.5065 | 1.912 | 0.03428 | 0.265 | 0.7913 | |
| SOA | 9.976 | 1.820 | 1.019e+04 | 5.483 | < 0.001 | *** |
| SFmask | 0.07532 | 0.621 | 1.019e+04 | 0.117 | 0.9066 | |
| StimulusType | 0.02686 | 0.01778 | 1.019e+04 | 1.511 | 0.1309 | |
| SOA:SFmask | -1.368 | 0.6421 | 1.019e+04 | -2.130 | 0.0332 | * |
| SFmask:StimulusType | -0.1022 | 0.01778 | 1.019e+04 | -5.747 | < 0.001 | *** |

Supplementary Table 7.

*OFA Fixed effects*

| | *Estimate* | *SE* | *df* | *T* | *p* | |
|---|---|---|---|---|---|---|
| (Intercept) | -13.73 | 3.9060 | 22.39 | -3.516 | 0.0019 | ** |
| SOA | 0.461 | 0.0174 | 10190 | 26.528 | < 0.001 | *** |
| SFmask | -3.722 | 1.7780 | 10190 | -2.093 | 0.0364 | * |
| StimulusType | 7.920 | 1.7780 | 10190 | 4.453 | < 0.001 | *** |
| SOA:SFmask | -0.0493 | 0.0174 | 10190 | -2.834 | 0.0046 | ** |
| SOA:StimulusType | -0.0660 | 0.0174 | 10190 | -3.799 | 0.0001 | *** |

### Supplementary material 5

#### LMM figure extra ROIs V2 and OFA

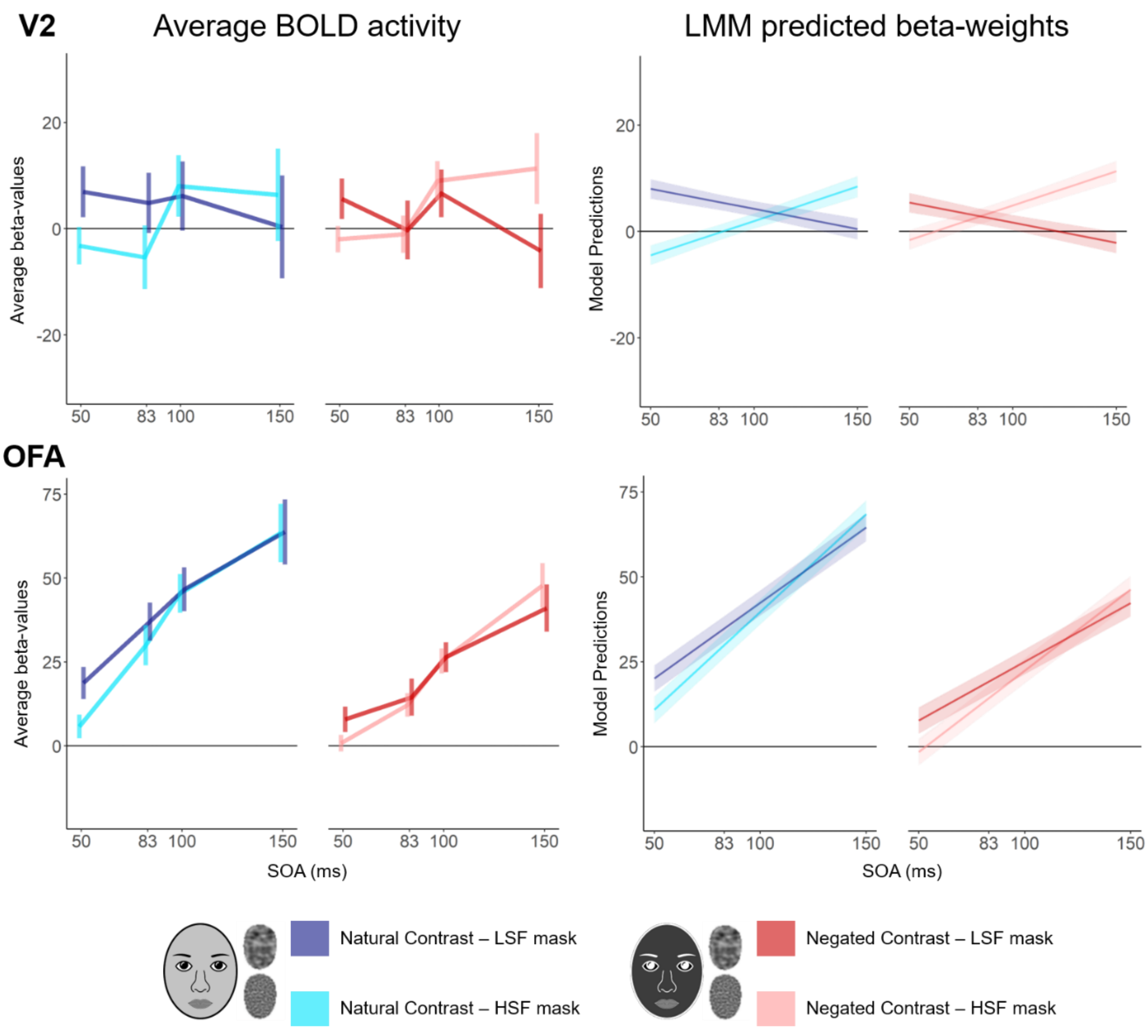


*Figure 3.* Intact-scrambled response differential for extra ROIs V2 (top; anatomically defined with Freesurfer label) and OFA (bottom; functionally defined with the functional localiser). Line plots in the left column represent the group averaged beta-values including a 95% within-subjects confidence interval (Loftus & Masson, 1994). The LMM predictions shown on the right plots are from a full model including all interaction effects with was ran just for illustrative purposes. Cartoon faces are used in replacement of photographs in the figure as journal regulations prohibit the use of identifiable faces.

#### LMM figure PPI analysis

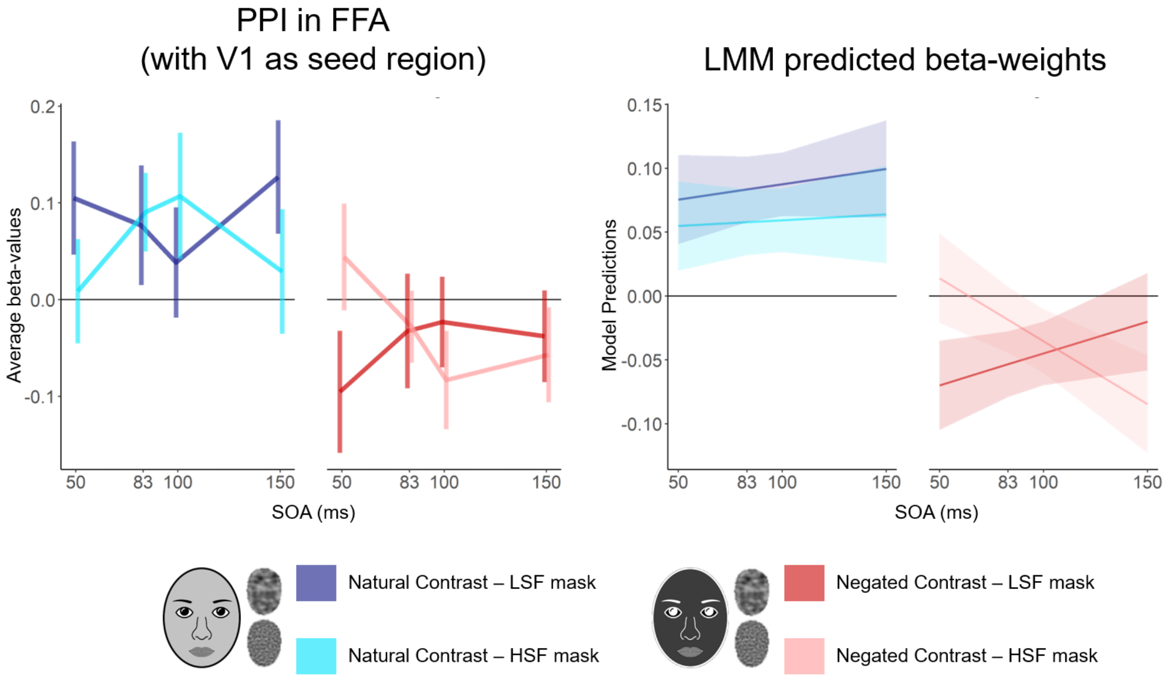


*Figure 4.* Results of the PPI analysis in the FFA with V1 as seed region. Left. Line plot shows PPI results averaged over all subjects including a 95% within-subjects confidence interval (Loftus & Masson, 1994). Right. Line plot with the model predictions of the full model including all interaction terms, since there were no significant three-way or two-way interactions (only a main effect of stimulus type), the plots are just for illustrative purposes. Cartoon faces are used in replacement of photographs in the figure as journal regulations prohibit the use of identifiable faces.
